# A peptide catalyst can replace an essential enzyme in a eukaryotic cell

**DOI:** 10.64898/2026.01.27.701630

**Authors:** Kira A. Podolsky, Oscar J. Molina, Vivian Y. Long, Ronald T. Raines

## Abstract

Protein enzymes are central to modern biology, yet how catalysis emerged before the evolution of large, folded proteins remains unresolved. Here we show that a short, genetically encoded peptide can replace an essential enzyme in a living eukaryotic cell. We designed minimal peptides containing a Cys-Xaa-Cys catalytic motif and an endoplasmic reticulum retention signal, and identified variants that rescue the otherwise lethal deletion of protein disulfide isomerase (PDI) in *Saccharomyces cerevisiae*. Cells relying on these peptides remain viable, though they grow more slowly and adapt by activating stress-response pathways, consistent with PDI being replaced by catalysts of lower intrinsic efficiency. Biochemical analyses show that peptide activity depends on local chemical environment and secondary structure rather than a globular fold. These results demonstrate that short peptides can replace an essential cellular reaction in vivo at the system level, supporting the plausibility of peptide-based catalysis as a precursor to modern protein enzymes.

## Main Text

Before the emergence of modern proteins, early life must have relied on simpler polymers to catalyze essential chemical reactions. Short peptides are attractive candidates for such primordial catalysts: they are chemically diverse, more stable than RNA, and readily generated by prebiotic chemistry (*1–5*). Yet in modern cells, catalysis is almost universally carried out by large, folded enzymes, and it remains unclear whether short peptides are sufficient to support essential biochemical processes in vivo. Demonstrating such activity in a living organism would establish a lower bound on the structural complexity required for biological catalysis.

Protein disulfide isomerase (PDI) provides a stringent test of this possibility. PDI is an abundant (∼0.2 mM) 56-kDa enzyme that catalyzes oxidative protein folding in the endoplasmic reticulum (ER) (*6*). Its deletion or inactivation is lethal in eukaryotes (*6–8*). Although nonnatural synthetic peptides (*9–11*) and even amino acids (*12*) can catalyze chemical reactions in vitro, no genetically encoded peptide has been shown to replace a full-length enzyme in vivo. Here, we ask whether a minimal peptide, localized to the ER and bearing a mimetic catalytic motif, can replace PDI in an organism.

### Minimal requirements for a catalyst of oxidative protein folding

PDI catalyzes oxidative protein folding in the ER by mediating thiol–disulfide exchange reactions using two CGHC active sites (Fig. 1A). In each, the thiol of the N-terminal cysteine residue has a depressed p*K*_a_ that promotes nucleophilic attack on nonnative disulfide bonds (*13*), initiating iterative rearrangement reactions that proceed until native structure is achieved (Fig. 1B) (*6, 14*). The C-terminal active-site cysteine has an elevated p*K*_a_ (*15*) and resolves unproductive mixed-disulfide intermediates, enabling catalyst turnover (*6*).

**Fig. 1.**
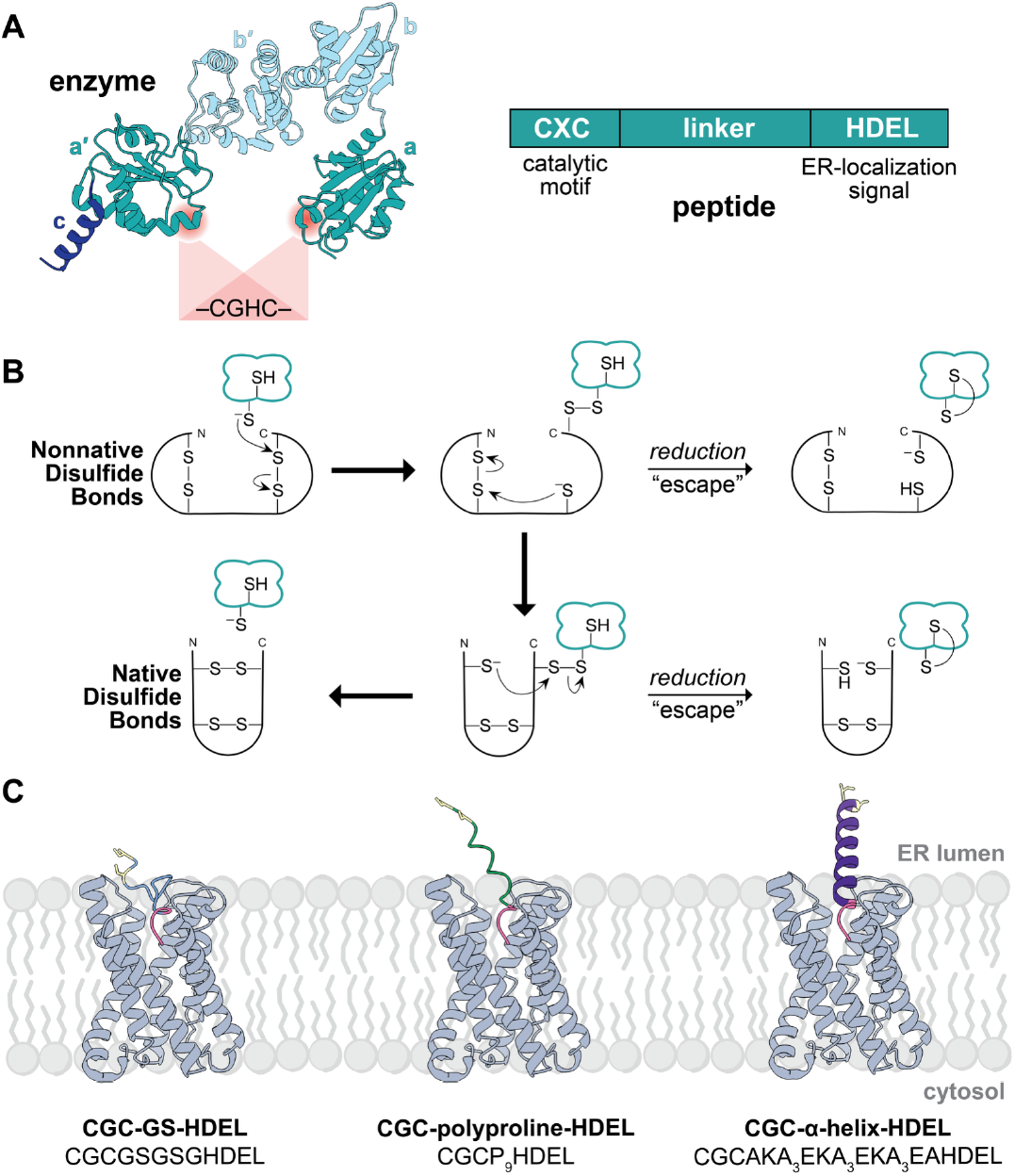
Structure and design principles underlying peptide replacement of protein disulfide isomerase. (**A**) Ribbon diagram of *S. cerevisiae* protein disulfide isomerase (PDI; PDB: 2b5e) showing its domains and two CGHC catalytic motifs, alongside a schematic representation of the minimal peptide catalysts used in this study. Replacement is operationally defined as the restoration of viability following genetic deletion of PDI by an alternative catalyst embedded within the cellular folding network. (**B**) Schematic of PDI-catalyzed disulfide bond isomerization. The N-terminal thiolate of the CGHC motif initiates thiol–disulfide exchange with nonnative disulfides, followed by iterative rearrangement (and, if necessary, resolution by the C-terminal cysteine), enabling catalytic turnover. (**C**) AlphaFold2-predicted models of three designed CXC-linker-HDEL peptides bound to the KDEL receptor (gray; PDB: 6i6h) within the ER membrane. Models were used to estimate linker length and orientation. Peptide sequences are shown below each model.

PDI shares ∼30% sequence identity with thioredoxin (Trx), a 12-kDa cytosolic oxidoreductase that also contains a CGPC active-site motif with a low thiol p*K*_a_ but lacks disulfide isomerase activity (*16–18*). Notably, yeast lacking *PDI1* (*pdi1*Δ) can be rescued by targeting *Escherichia coli* Trx to the ER and substituting its active site with the CGHC motif of PDI (*19*). This functional substitution correlates with redox tuning: the CGHC motif of PDI has a disulfide reduction potential (*E*°′ = −0.180 V) that matches the ER environment, whereas wild-type Trx is substantially more reducing (*E*°′ = −0.270 V) (*19–21*). These features—low thiol p*K*_a_ and high disulfide *E*°′—are the minimal chemical requirements for a catalyst of oxidative protein folding (*22*).

In CX_*n*_C peptides, the disulfide *E*°′ depends strongly on the number of intervening residues (*23*). A minimal H-Cys-Gly-Cys-NH_2_ peptide exhibits a disulfide reduction potential (*E*°′ = −0.167 V) that closely matches that of PDI (*24*), along with a low thiol p*K*_a_ value because of favorable Coulombic interactions of the thiolate and N-terminal ammonium group (*25*). Consistent with these attributes, this tripeptide catalyzes disulfide bond formation, cleavage, and rearrangement in vitro, functionally mimicking PDI despite lacking a globular fold (*24, 26*). These observations led us to hypothesize that simple peptides bearing a CXC catalytic motif, when localized to the ER, could be sufficient to support the viability of an organism.

### Design of minimal ER-localized peptide catalysts

We designed minimal peptides containing a CXC catalytic motif and an ER retention signal to test whether such short sequences could substitute for PDI in vivo (Fig. 1A). Each peptide consists of: (1) an N-terminal CXC motif, (2) a C-terminal HDEL sequence for ER retention, and (3) a linker region to present the catalytic motif to client proteins. We refer to these constructs as “CXC-linker-HDEL” peptides.

We reasoned that an effective linker would mimic the spatial separation between the activesite of Trx and the HDEL receptor in constructs that replace PDI (*19*). AlphaFold3 predictions estimated this distance to be ∼28 Å (Figs. S1 and S2; Table S1). Based on this distance, we designed three linkers (Fig. 1C): a flexible GSGSG motif (GS), a rigid polyproline stretch (P_9_) (*27*), and a stabilized α-helical segment (AKA_3_EKA_3_EKA_3_EA) (*28*). The resulting peptides, CXC-GS-HDEL (12 residues), CXC-polyproline-HDEL (16 residues), and CXC-α-helix-HDEL (24 residues), have calculated linker lengths of 25 Å, 37 Å, and 30 Å, respectively (Table S2). To validate these structural predictions, we synthesized the corresponding CGC-linker-HDEL peptides (Tables S3–S5) and confirmed their secondary structure using circular dichroism spectroscopy (Fig. 2A).

**Fig. 2.**
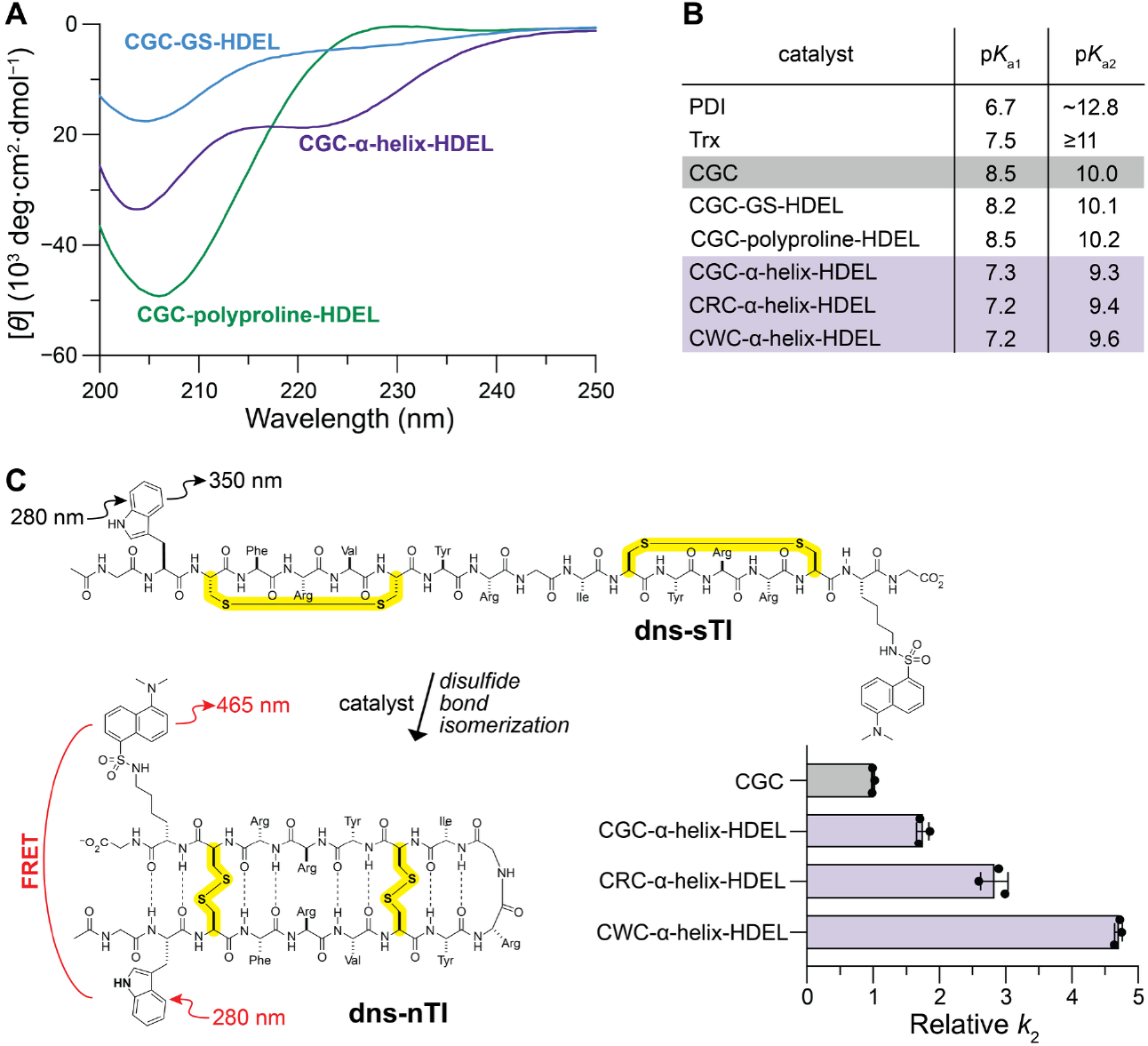
Structural and catalytic properties of synthetic peptide catalysts. (**A**) Circular dichroism spectra of CGC-linker-HDEL peptides in phosphate-buffered saline. CGC-GS-HDEL exhibits a random-coil spectrum (blue), CGC-polyproline-HDEL shows features characteristic of a polyproline II helix (green), and CGC-α-helix-HDEL displays α-helical minima at ∼208 and 222 nm (purple). (**B**) p*K*_a_ values of the active site N-terminal (p*K*_a1_) and C-terminal (p*K*_a2_) cysteine residues of PDI (*13, 15*), Trx (*18*), and CXC peptides, illustrating that context can tune cysteine reactivity. (**C**) Schematic of the fluorescence resonance energy transfer (FRET) assay used to measure disulfide bond isomerization. Conversion of scrambled tachyplesin I (dns-sTI) to its native disulfide form (dns-nTI) increases FRET emission at 465 nm upon excitation at 280 nm. *Inset*: Graph of *k*_2_ (± SE) for CXC-α-helix-HDEL peptides relative to the CGC peptide.

### Replacing PDI with a peptide catalyst in vivo

To test whether minimal CXC-linker-HDEL peptides can substitute for PDI, we used plasmid shuffling (*29*) in *S. cerevisiae* (Fig. 3A). Because the optimal “X” residue was unknown, we generated a random library covering all 20 canonical amino acids and the three different linkers, yielding 60 constructs (Table S6).

**Fig. 3.**
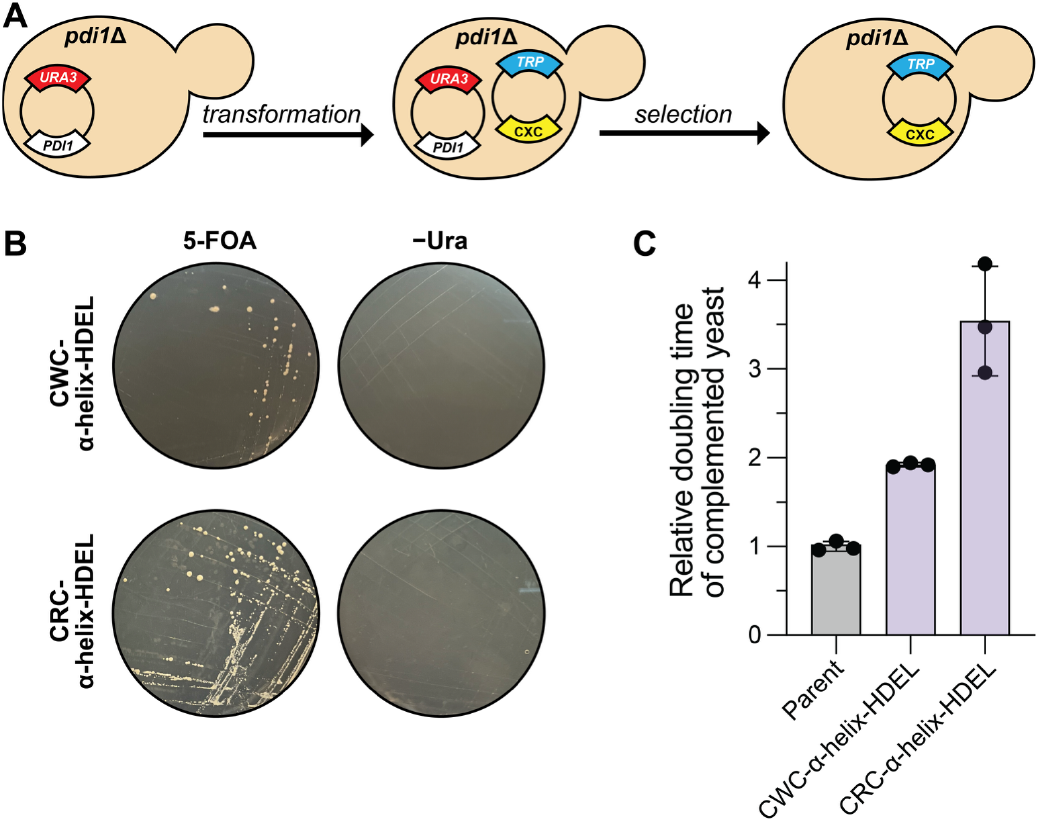
Genetic replacement of PDI by peptide catalysts in yeast. (**A**) Schematic of plasmid shuffling used to test peptide-mediated replacement of PDI. A *pdi1*Δ strain initially carrying a *URA3*-marked PDI plasmid is transformed with a *TRP1*-marked plasmid encoding a CXC-linker-HDEL peptide. Iterative selection on −Trp and 5-fluoroorotic acid (5-FOA) yields strains dependent on the peptide catalyst. (**B**) *pdi1*Δ yeast producing CWC-α-helix-HDEL or CRC-α-helix-HDEL grow on 5-FOA plates but not −Ura plates, demonstrating that the peptide catalysts replace PDI as the sole source of oxidative protein folding activity required for viability. (**C**) Doubling times (± SE) of peptide-replaced yeast strains relative to the parent strain, reflecting establishment of viable but lower-efficiency cellular states following enzyme replacement.

We began with a *pdi1*Δ yeast strain carrying plasmid pCT37, which expresses PDI and the negative selectable marker *URA3*. These cells were transformed with plasmid YEpWL.CXClinkerHDEL, which expresses the library of CXC-linker-HDEL peptides fused to an α-factor leader sequence (*30*) and a positive selectable marker, *TRP1*. The transformed cells were grown on solid synthetic dropout medium lacking tryptophan (−Trp). Colonies formed within seven days and were replica plated onto medium containing 5-fluoroorotic acid (5-FOA), which selects against *URA3* on pCT37 (*31*). Cells that survive this selection indicate successful complementation by the peptide.

The cycle of −Trp and 5-FOA selection was repeated four times. Colony growth was also monitored on −Ura medium to confirm loss of plasmid pCT37. Successful complementation produces growth on 5-FOA but not on −Ura, reflecting dependence on the peptide rather than PDI. Negative controls, including transformations with no plasmid or with an unrelated enzyme, RNase A, failed to grow on 5-FOA or −Trp, confirming that *pdi1*Δ yeast cannot survive without PDI or a functional peptide (Fig. S3). A positive control demonstrated that yeast PDI restores viability after four selection cycles (Fig. S4).

After four rounds of selection, complementing clones were observed exclusively in the α-helix linker library, but not in the GS or polyproline libraries (Fig. S5). Growth on 5-FOA medium and the absence of growth on −Ura confirmed that these peptides replace PDI in sustaining viability (Fig. 2B).

### Function of catalytic peptides in vivo

Complemented yeast strains were genotyped by PCR (Fig. S6; Tables S7 and S8), and gel-extracted products were sequenced. Two peptides, CWC-α-helix-HDEL and CRC-α-helix-HDEL, were identified as complementing catalysts (Fig. S7). Microscopy revealed that the morphology of complemented cells resembled that of the parental *S. cerevisiae* strain (Fig. S8).

To assess catalytic function in vivo, we measured growth rates of complemented *pdi1*Δ cells. The doubling times of cells expressing CWC-α-helix-HDEL and CRC-α-helix-HDEL were only approximately 2.0- and 3.5-fold greater than that of the parent strain, respectively (Fig. 3C). These results indicate that both peptides can substitute for PDI in vivo, replacing its essential catalytic function, albeit with reduced efficiency.

### Biochemical attributes of catalytic peptides

We next characterized key biochemical properties of the CWC-α-helix-HDEL and CRC-α-helix-HDEL peptides. Key CXC and CXC-linker-HDEL constructs were synthesized (Tables S3–S5). The N-terminal cysteine of CXC peptides with an α-helix linker has a p*K*_a_ near 7, comparable to PDI (Figs. 2B and S9), whereas CGC, CGC-GS-HDEL, and CGC-polyproline-HDEL constructs have higher p*K*_a_ values near 8.5. This shift likely arises from the α-helix dipole, which withdraws electrons from the N-terminal cysteine, increasing the fraction of thiolate (*32*). The identity of the “X” residue had minimal effect on p*K*_a_ within CXC-linker-HDEL peptides.

We then measured disulfide isomerase activity in vitro using a fluorescence resonance energy transfer (FRET) assay (*26*). The substrate was a modified peptide derived from tachyplesin I (TI), a β-hairpin stabilized by two native disulfides. A scrambled version of TI was synthesized with pendant tryptophan and dansyllysine residues (dns-sTI), such that isomerization to the native disulfide form (dns-nTI) brings the fluorophores into proximity, enabling FRET (Fig. 2C).

We found that the ability of peptides to mediate the isomerization of the dns-sTI substrate depended on the X residue, following the trend: CGC < CGC-α-helix-HDEL < CRC-α-helix-HDEL < CWC-α-helix-HDEL (Fig. 2C, inset). The tryptophan residue in CWC-α-helix-HDEL nearly doubled the activity relative to arginine in CRC-α-helix-HDEL, mirroring the difference in doubling times of complemented cells (Fig. 3C) and linking chemical reactivity to in vivo viability.

### Transcriptomics of peptide-complemented cells

Next, we explored why cells relying on peptide surrogates for PDI grow more slowly than the parental strain (Fig. 3C). We hypothesized that replacing PDI with catalysts of lower intrinsic efficiency triggers stress-response and compensatory pathways. To test this hypothesis, we used RNA sequencing (RNA-seq) (*33*) to identify differentially expressed genes (DEGs) between peptide-complemented and parental yeast. Gene ontology (GO) analysis classified DEGs into biological processes that differed between the strains.

*S. cerevisiae* manages ER stress primarily through the unfolded protein response (UPR) and ER-associated degradation (ERAD) pathways (*34, 35*). The UPR upregulates chaperones to maintain ER folding homeostasis, whereas ERAD targets misfolded proteins for degradation.

GO analysis revealed broad transcriptional changes in peptide-complemented cells. Ribosome biogenesis and cytosolic translation were generally downregulated, whereas mitochondrial translation was upregulated (Fig. 4A; Tables S9 and S10). Notably, CWC-α-helix-HDEL cells, but not CRC-α-helix-HDEL cells, showed significant downregulation of ribosome biogenesis and cytosolic translation, which can reduce the accumulation of misfolded proteins. Upregulation of mitochondrial translation in both strains likely reflects an increased energy demand under oxidative stress, consistent with mitochondrial enlargement observed during the UPR and diauxic shifts (*36*).

**Fig. 4.**
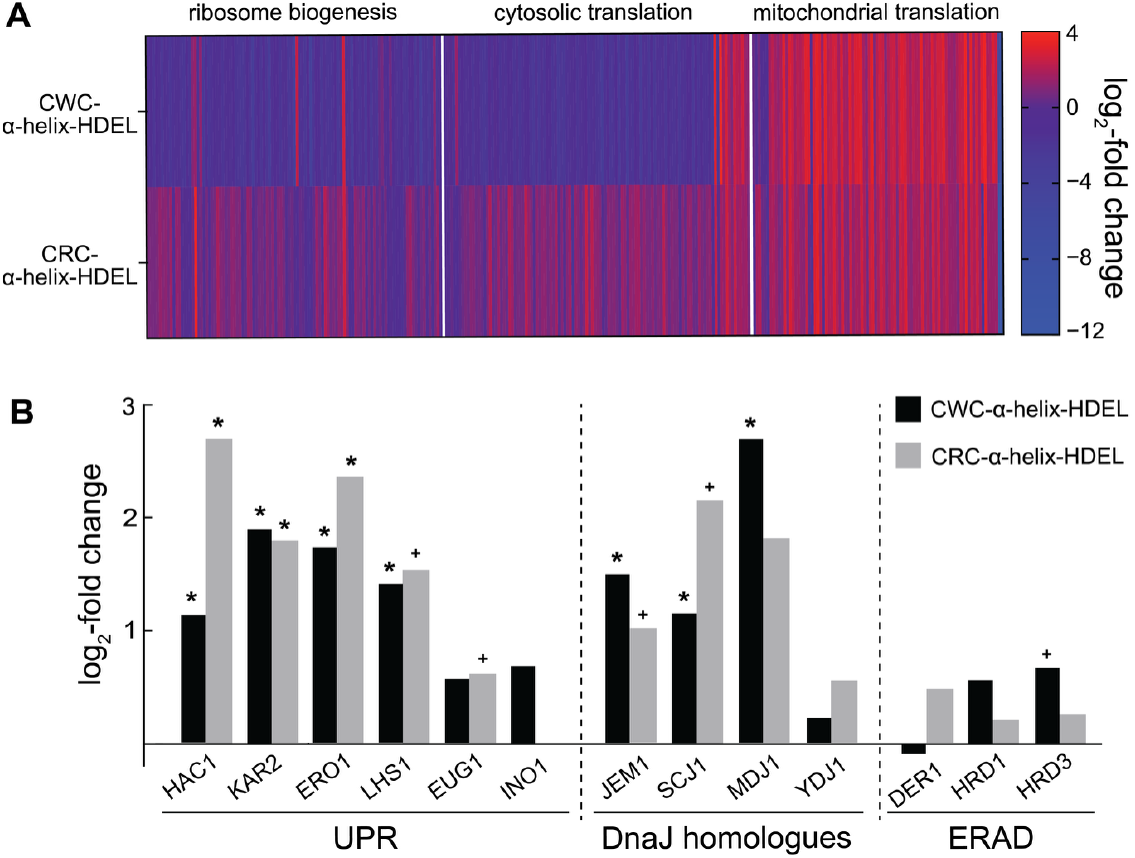
Cellular network remodeling following replacement of PDI with a peptide catalyst. (**A**) log_2_-fold changes in transcript abundance for genes associated with ribosome biogenesis, cytosolic translation, and mitochondrial translation in *pdi1*Δ yeast producing CWC-α-helix-HDEL or CRC-α-helix–HDEL yeast relative to the parent strain. Each vertical line represents an individual gene; color indicates direction and magnitude of change, reflecting reprogramming of translational and metabolic pathways following enzyme replacement. (**B**) Differential expression of unfolded protein response (UPR), DnaJ-family chaperone, and endoplasmic reticulum– associated degradation (ERAD) genes in peptide-replaced yeast relative to the parent strain. Activation of UPR and selective chaperone pathways indicates compensatory network remodeling that enables viability following replacement of PDI. * denotes *P*_adj_ < 0.05; ^+^ denotes *P* < 0.05.

To examine UPR activation, we assessed key components of the pathway. Misfolded protein accumulation in the ER triggers dissociation of the chaperone KAR2 from the transmembrane sensor Ire1p, resulting in noncanonical splicing of HAC1 mRNA (*37*). The Hac1p transcription factor then upregulates UPR elements (UPREs). DEG analysis confirmed that both CWC-α-helix-HDEL and CRC-α-helix-HDEL cells significantly upregulate UPR-related genes, including HAC1, KAR2, LHS1 (an Hsp70 chaperone), and ERO1 (Fig. 4B; Table S11) (*38, 39*). Interestingly, EUG1, a nonessential ER disulfide isomerase, was not upregulated, indicating reliance on the peptide catalyst in conjunction with ERO1. INO1 expression remained unchanged, suggesting steady ER membrane synthesis.

In contrast to UPR activation, transcripts associated with the ERAD pathway were not significantly affected, indicating that misfolded proteins are not being preferentially degraded. Instead, select chaperones of the DnaJ family were upregulated. ER-resident homologues JEM1 and SCJ1, along with the mitochondrial homologue MDJ1, were significantly upregulated, consistent with increased mitochondrial activity, whereas the cytosolic homologue YDJ1 remained unchanged (*40*).

Overall, transcriptomic analysis shows that replacing an essential enzyme with a peptide catalyst drives adaptive remodeling of the cellular folding network, establishing a new homeostatic state compatible with viability.

## Discussion and Outlook

We have identified peptide catalysts capable of replacing an essential enzyme in a eukaryotic cell. Through rational design and genetic selection, CRC-α-helix-HDEL and CWC-α-helix-HDEL were found to complement *pdi1*Δ *S. cerevisiae*. In vitro characterization suggests that these peptides are effective because of the chemical environment provided by the α-helix and the specific residues within the CXC motif. The α-helical linker lowers the p*K*_a_ of the N-terminal cysteine, increasing the fraction of reactive thiolate. The tryptophan residue in the CWC motif could enhance catalysis by engaging misfolded proteins through the hydrophobic effect (*41*), while the arginine residue in the CRC motif could interact with anionic proteins, which are especially abundant in the ER lumen (*42*).

In a cell, PDI does not function as an autonomous catalyst, nor do the complementing peptides operate independently of the cellular environment. In both cases, viability emerges from reciprocal interactions between the catalyst and the folding, quality-control, and stress-response networks of the cell. Consistent with this system-level mode of replacement, peptide-dependent cells exhibit slower growth and activate the unfolded protein response, reflecting the cellular burden imposed by catalysts with lower intrinsic efficiency (*43*). This codependence between catalytic activity and network remodeling may reflect an ancestral mode of catalysis, in which modestly active chemical motifs acted in concert with their environments before the evolution of folded protein scaffolds. In this sense, the peptides fully replace PDI as system-level catalysts of oxidative protein folding.

Our findings provide insight into how short peptides could have served as primitive catalysts in early life. If such peptides were once sufficient to sustain essential chemistry, remnants of this mode of catalysis could persist today among the many intracellular and extracellular peptides of unknown function (*44, 45*). Peptides with modest catalytic activity may have facilitated essential reactions at slower rates than modern enzymes, with secondary structure and specific residues enhancing function. Over evolutionary time, such peptide-based catalytic motifs could have been elaborated through sequence expansion and structural stabilization, ultimately giving rise to folded enzymes with higher efficiency and broader substrate scope. More broadly, our results highlight the critical roles of both primary sequence and secondary structure in determining catalytic activity, offering a mechanistic framework for understanding the emergence of enzymatic function from short peptides.

## Acknowledgments

We are grateful to Prof. Jakob R. Winther (University of Copenhagen) for advice on yeast strains. We thank Dr. Ky Lowenhaupt (MIT Biophysical Instrumentation Facility) for assistance with circular dichroism spectroscopy and Dr. Mohan Kumar (MIT Department of Chemistry Instrumentation Facility) for assistance with mass spectrometry.

## Funding

Schmidt Science Fellows, in partnership with the Rhodes Trust (KAP) National Institutes of Health Grant R35 GM148220 (RTR)

## Author contributions

Conceptualization: KAP, RTR

Methodology: KAP, RTR

Investigation: KAP, OJM, VYL

Funding acquisition: KAP, RTR

Supervision: KAP, RTR

Writing—original draft: KAP, OJM, VYL

Writing—review & editing: KAP, OJM, VYL, RTR

## Competing interests

Authors declare that they have no competing interests.

## Data and materials availability

All data are available in the main text or the supplementary materials.

## Notes

### Competing Interest Statement

The authors have declared no competing interest.

### Summary of Updates

Mislabeling of cartoon images in Figs. 1C and S2, and the addition of a plasmid map to the Supplementary Materials.

