## Supplementary Materials for "A peptide catalyst can replace an essential enzyme in a eukaryotic cell"

**The PDF file includes:**

Materials and Methods
Figs. S1 to S9
Tables S1 to S11
References (*1–16*)

### Abbreviations

| | |
|---|---|
| ACN | acetonitrile |
| Arg or R | (2*S*)-arginine |
| bp | basepair |
| cDNA | complementary DNA |
| Cys or C | (2*S*)-cysteine |
| DCM | dichloromethane |
| DEG | differentially expressed gene |
| DIC | diisopropylcarbodiimide |
| DIEA | *N*,*N*-diisopropylethylamine |
| $DiOC_6$ | 3,3′-dihexyloxacarbocyanine iodide |
| DMF | dimethylformamide |
| dns | dansyl or 5-(dimethylamino)naphthalene-1-sulfonyl |
| DODT | 2,2′-(ethylenedioxy)diethanethiol |
| dUTP | 2′-deoxyuridine-5′-triphosphate |
| EDTA | ethylenediaminetetraacetic acid |
| ER | endoplasmic reticulum |
| ERAD | endoplasmic reticulum-associated degradation |
| ESI | electrospray ionization |
| Fmoc | 9-fluorenylmethyloxycarbonyl |
| FRET | fluorescence resonance energy transfer |
| Glu or E | (2*S*)-glutamic acid |
| Gly or G | glycine |
| GO | gene ontology |
| HPLC | high-performance liquid chromatography |
| His or H | (2*S*)-histidine |
| LC–MS | liquid chromatography–mass spectrometry |
| Leu or L | (2*S*)-leucine |
| Lys or K | (2*S*)-lysine |
| nTI | native tachyplesin I |
| OD | optical density |
| Oxyma | ethyl cyano(hydroxyimino)acetate |
| PAE | predicted alignment error |
| PBS | phosphate-buffered saline |
| PCR | polymerase chain reaction |
| PDI | protein disulfide isomerase (EC 5.3.4.1; *Saccharomyces cerevisiae*: UniProtKB P17967) |
| pLLDT | predicted local distance difference test |
| Pro or P | (2*S*)-proline |
| $P_{adj}$-value | adjusted probability value |
| $P$-value | probability value |
| rcf | relative centrifugal force |
| RNase A | ribonuclease A (EC 4.6.1.18; *Bos taurus*: UniProtKB P61823) |
| RNA-seq | RNA-sequencing |
| SE | standard error |
| Ser or S | (2*S*)-serine |

| | |
|---|---|
| sTI | disulfide-scrambled tachyplesin I |
| TFA | trifluoroacetic acid |
| TI | tachyplesin I |
| TIPS | triisopropylsilane |
| Tris | tris(hydroxymethyl)aminomethane |
| Trp or W | (2*S*)-tryptophan |
| Trt | trityl or triphenylmethyl |
| Trx | thioredoxin (*Escherichia coli*: UniProt P0AA25) |
| UPR | unfolded protein response |
| UPRE | unfolded protein response element |
| Xaa or X | any α-amino acid residue |
| YPD | yeast peptone dextrose |

### Materials and Methods

#### Yeast strains, plasmids, and reagents

The *Saccharomyces cerevisiae* parent strain W303-1Bα Δ*pdi1*/pCT37 (*ade2-1 can1-100 ura3-1 leu2-3,112 trp1-1 his3-11,15 pdi1::HIS3*) was a generous gift from Carlsberg A/S (Copenhagen, Denmark). Plasmid pCT37 (*S. cerevisiae PDI1* and *URA3*) was a generous gift from Prof. Tom H. Stevens (University of Oregon) (*1*). Plasmids YEpWL.TRX (*TRX* and *TRP1*), pRS424 (*S.cerevisiae PDI1* and *TRP1*), and YEpWL.RNaseA (*RNaseA* and *TRP1*) were available from our previous studies (*2*, *3*).

α-Chymotrypsin was purchased from Sigma–Aldrich (St. Louis, MO, USA) 5-FOA was purchased from Zymo Research (Irvine, CA, USA). Synthetic dropout −Trp media was purchased from Takara (Kusatsu, Shiga, JP). Synthetic dropout −Ura media was purchased from Sigma–Aldrich. milli-Q water was obtained from a Milli-Q Q-Pod water purification system from Millipore Sigma (Burlington, MA, USA).

All peptide synthesis reagents were purchased from Iris Biotech (Marktredwitz, Germany), CEM (Matthews, NC, USA), Sigma−Aldrich, or TCI (Portland, OR, USA). Fmoc-Lys(dansyl)-OH was purchased from ChemPep (Wellington, FL, USA). Trifluoroacetic acid (TFA), triisopropylsilane (TIPS), and 2,2′-(ethylenedioxy)diethanethiol (DODT) were purchased from Sigma–Aldrich.

#### AlphaFold predictions

Structures of CXC-linker-HDEL peptides were predicted using ColabFold (*4*). Multiple sequence alignments were constructed using the MMseq2 against the environmental and UniRef databases of ColabFold. Default ColabFold parameters were used, with five independent models per sequence and three recycles. The resulting structures were all energy-minimized via Amber relaxation. Predicted structure confidence was determined using the per-residue predicted local distance difference test (pLLDT) score and the predicted aligned error (PAE) matrices. A pLDDT threshold >70 is considered a high-confidence prediction for local backbone structure and geometry. Structural models were visualized using UCSF ChimeraX version 1.6.1. Amino acid sequences for the peptides are listed in table S2.

Structures of CXC-linker-HDEL peptides bound to the human KDEL receptor were predicted using ColabFold (*4*). Sequences were separated by ":" to specify inter-protein chain breaks for modeling complexes. Multiple sequence alignments were constructed using the

MMseq2 against the environmental and UniRef databases of ColabFold. Default ColabFold parameters were used, with five independent models for each sequence and three recycles. The resulting structures were all energy-minimized via Amber relaxation. Structural models were visualized using ChimeraX. The structure of wild-type thioredoxin with a C-terminal HDEL bound to the human KDEL receptor was predicted using AlphaFold Server and "1" as the seed number (*5*). Amino acid sequences for each protein·protein or peptide·protein complex structure are listed in table S1.

#### Estimation of linker lengths

Thioredoxin variants can replace *S. cerevisiae* protein disulfide isomerase (PDI) when localized to ER receptors via a C-terminal HDEL motif (*2*). The distance between the HDEL motif and the catalytic CGPC motif in thioredoxin is the minimal distance known to complement PDI in yeast. We generated models of wild-type thioredoxin with a C-terminal HDEL motif bound to the human KDEL receptor (described above). Using these models (fig. S1), the distance (28 Å) between $C^{\alpha}$ of Cys32 (which is the N-terminal cysteine residue in the CGPC motif) and $C^{\alpha}$ of Val105 (which is conjugated to the HDEL motif) was measured using UCSF ChimeraX version 1.6.1. CXC-linkers were designed to span this distance. The estimated lengths of each CXC-linker were calculated using the average length of an amino acid residue (for the GS linker), the rise per residue in a polyproline type-II helix (for the polyproline linker), and the rise per residue in an α-helix (for the α-helix linker) (table S2).

#### Microwave-assisted solid-phase peptide synthesis

The CGC and CXC-linker-HDEL peptides were synthesized via SPPS on a Liberty Blue peptide synthesizer from CEM (Matthews, NC, USA) on a 0.1 mmol synthesis scale (table S2). The Fmoc-protected amino acids (0.2 M in DMF), coupling agent (20% v/v solution of 4-methylpiperidine in DMF), wash solvent (DMF), activator (0.5 M DIC in DMF), and base activator (0.5 M Oxyma and 0.1 equiv DIEA in DMF) were pre-loaded onto the instrument before microwave-assisted synthesis. Either 0.182 g of Fmoc-Cys(Trt)-Rink resin (0.22 mmol/g loading) for CGC or 0.455 g of Fmoc-Leu-Wang resin (0.55 mmol/g loading) for the CXC-linker-HDEL peptides was added to the 30 mL Liberty Blue reaction vessel. During synthesis, amino acids were coupled under standard microwave-assisted deprotection and coupling settings. Post-synthesis, the resin was removed from the reaction vessel, washed with 5 mL of DCM, and left overnight in a desiccator for further deprotection and purification.

#### Deprotection and purification of catalytic peptides

The peptide-conjugated resin was incubated on a rotator for 45 min in a 5 mL TFA/TIPS/water/DODT deprotection cocktail (36.8:1:1:1). The filtrate was collected into two 15 mL Falcon tubes and precipitated with 14 mL of ice-cold diethyl ether. The tubes were subjected to centrifugation at 3460 rcf for 4 min at 4 °C, and the supernatant was discarded. This process was repeated twice with 7 mL of ice-cold diethyl ether to resuspend the pellet. After the final diethyl ether wash, the pellet in each tube was dissolved in 1 mL of ACN and 3 mL of milli-Q water, frozen at −80 °C, and lyophilized overnight. The lyophilized crude peptide powder was dissolved in milli-Q water for HPLC purification (table S4). Purification was done for CGC using a Vydac protein and peptide C18 semipreparatory column and for CXC-linker-HDEL peptides using an XSelect Peptide C18 preparatory column. Purification used linear Phase A/Phase B gradients (Phase A: 95% v/v $H_2O$ + 5% v/v ACN + 0.1% v/v TFA; Phase B: 95% v/v ACN + 5% v/v $H_2O$ + 0.1% v/v TFA) on a 1260 Infinity II HPLC instrument from

Agilent Technologies (Lexington, MA, USA). The pure fractions were pooled, frozen at −80 °C, and lyophilized overnight. Purified peptides, obtained as a white powder, were analyzed by quadrupole time-of-flight (Q-TOF) liquid chromatography–mass spectrometry (LC–MS) in electrospray ionization positive mode using a Poroshell 120 EC-C18 column on a 6530C mass spectrometer or a 6125B mass spectrometer equipped with an Agilent 1260 Infinity LC from Agilent Technologies (Santa Clara, CA, USA) (table S5). Purified peptides were stored under $N_2$(g) in a −20 °C freezer.

**Determination of peptide concentration with Ellman's assay**

The sulfhydryl concentration of the CGC and CXC-linker-HDEL peptides was calculated via Ellman's assay (*6*). Ellman's reagent solution (4 mg/mL) was prepared in Ellman's reaction buffer (which was 100 mM sodium phosphate buffer, pH 8.0, containing 1.0 mM EDTA). Tubes for each peptide and blank were prepared with 50 µL of Ellman's reagent solution and 2.75 mL of Ellman's reaction buffer. A 50 µL solution of peptide (or 50 µL of Ellman's reaction buffer for the blank) was added to the tubes. The solution was incubated for 15 min, after which the absorbance was measured at 412 nm on a Cary 60 UV–vis spectrophotometer from Agilent Technologies. The sulfhydryl concentration was calculated using the Beer–Lambert law:

$$c = \frac{A}{l\varepsilon}$$

In this equation, $c$ = concentration, $A$ = absorbance, $l$ = path length, and $\varepsilon$ = molar absorptivity, which is $\varepsilon$ = 14,150 $M^{-1}$ $cm^{-1}$ at 412 nm for 2-nitro-5-thiobenzoate (*6*).

The calculated sulfhydryl concentrations of the peptides were divided by two to account for the two sulfhydryl groups and used as the concentration of reduced peptides in experiments.

**Circular dichroism spectroscopy**

Lyophilized peptides (CGC-GS-HDEL, CGC-polyproline-HDEL, and GTG-α-helix-HDEL) were reconstituted to 0.25 mg/mL in phosphate-buffered saline (PBS), pH 7.2, at 25 °C. Samples and a PBS blank were run in triplicate on a J-1500 circular dichroism spectrophotometer from Jasco (Oklahoma City, OK, USA) at the MIT Biophysical Instrumentation Facility. All samples and blanks were averaged, and the data were baseline-corrected by subtracting the blank average from the peptide-containing sample averages.

**Plasmid library preparation**

The construction of the genetic library transgene used to insert into the YEpWL.Trx plasmid backbone was prepared by a Splicing-by-Oligo-(Overlap)-Extension PCR as described previously (*7*). The library transgenes encode CXC-GS-HDEL, CXC-polyproline-HDEL, and CXC-α-helix-HDEL peptides (table S6).

Briefly, oligonucleotides containing 40–60 nucleotides with overlap to each other were purchased from IDT. One codon position, encoding "X" in CXC, was created using an NNB strategy. The NNB library represents all 20 canonical amino acids, using 48 of 64 possible codons, while avoiding the incorporation of two stop codons (*8*). Q5 High-Fidelity DNA polymerase from New England Biolabs (Ipswich, MA, USA) was used to amplify the synthetic transgene with 5′ and 3′ flanking primers containing *Acc*65I and *Sal*I restriction enzyme recognition sites. The final PCR product was expected to be ~200 base pairs. The PCR reaction products were separated on a 2% w/v agarose gel. After separation, the amplicon was gel-extracted and purified using the Monarch DNA Gel Extraction kit from New England Biolabs.

The purified amplicon was then digested with restriction enzymes *Acc*65I and *Sal*I from New England Biolabs. In parallel, the YEpWL.Trx plasmid was digested with *Acc*65I and *Sal*I and then dephosphorylated using Quick CIP from New England Biolabs. The digested and treated backbone was then purified using a Monarch Spin PCR & DNA Cleanup Kit from New England Biolabs. The digested amplicon and the digested plasmid were incubated together according to the protocol for the Instant Sticky-end Ligase Master Mix from New England Biolabs, resulting in YEpWL.CXClinkerHDEL plasmid products.

The mixture of plasmids was transformed into 10-beta chemically competent, high-efficiency *E. coli* cells from New England Biolabs, and the cells were grown overnight at 37 °C with shaking at 250 rpm in Luria–Bertani (LB) medium containing 200 mg/mL ampicillin from Sigma–Aldrich. LB medium was prepared with 10 g of tryptone from Research Products International (Mount Prospect, IL, USA), 10 g of sodium chloride from Thermo Fisher Scientific (Waltham, MA, USA), and 5 g of yeast extract tryptone from Research Products International in 950 mL of milli-Q water and mixed until all solids were dissolved. The solution was adjusted to pH 7.9 using aqueous sodium hydroxide. The solution was adjusted to a final volume of 1.00 L with milli-Q water. This liter was autoclaved at 121 °C for 15 min and stored at room temperature. After 16–20 hours of *E. coli* outgrowth, a Monarch Spin Plasmid Miniprep Kit from

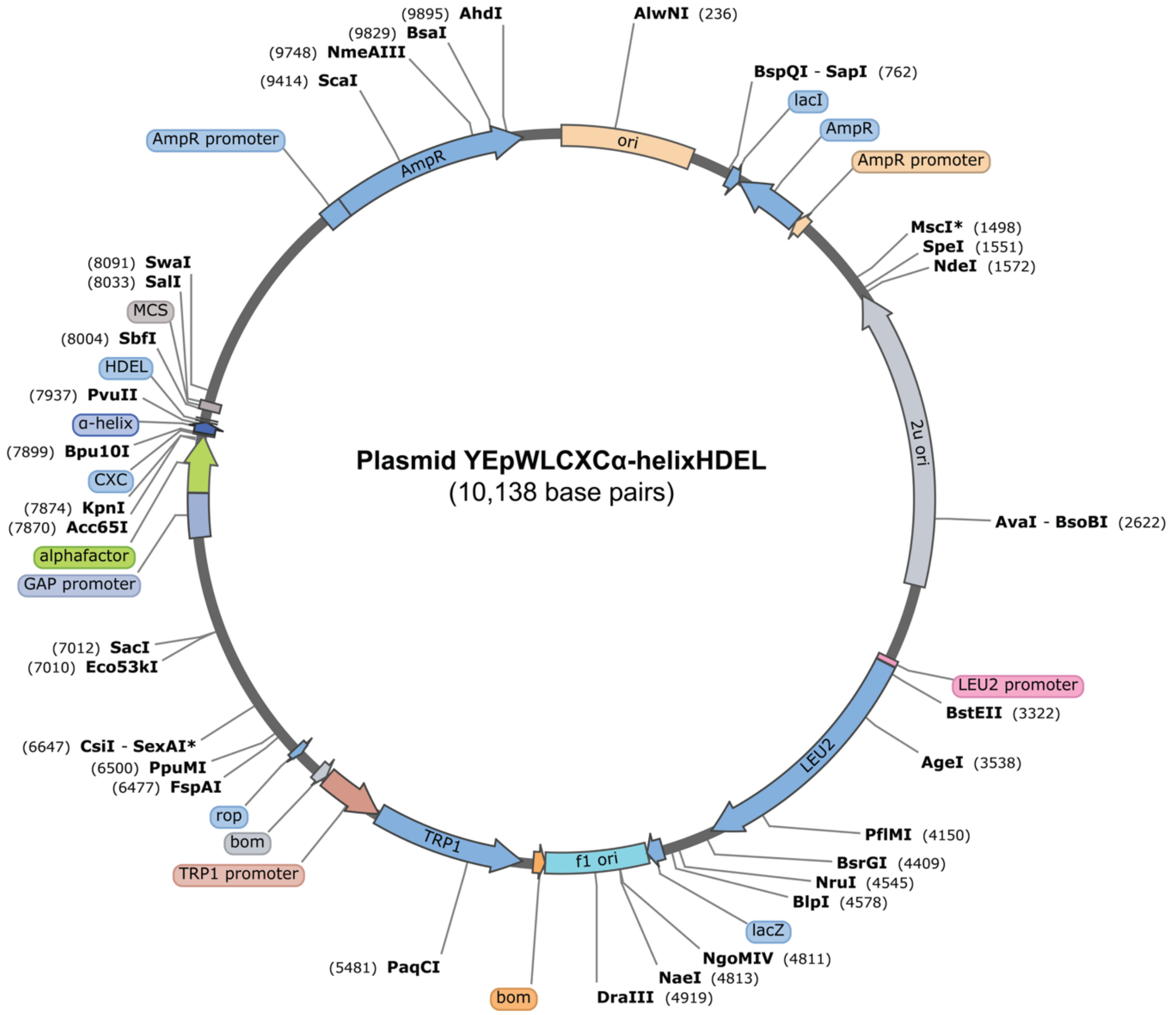

New England Biolabs was used to extract the YEpWL.CXClinkerHDEL plasmids, which were then sent to Plasmidsurus (San Francisco, CA, USA) for sequencing. Sequences were analyzed with SnapGene v6.0 software from GSL Biotech (Boston, MA, USA).

#### Yeast cell culture

Non-selective yeast cell culture was in yeast peptone dextrose (YPD) media. YPD was prepared by autoclaving 20 g of bacteriological peptone from VWR and 10 g of bacteriological yeast extract from Research Products International in 1.00 L of milli-Q water and autoclaving at 121 °C for 15 min. Then, 40 mL of 0.22 μm syringe-filtered 50% (w/v) glucose from Sigma–Aldrich was added to the cooled media. −Trp media was prepared by autoclaving one pouch of synthetic dropout −Trp powder from Takara (Kusatsu, Japan) in 0.50 L of water. −Ura media was prepared by autoclaving 0.96 g of −Ura dropout supplement and 3.35 g of yeast nitrogen base (without amino acids and with ammonia) in 500 mL of milli-Q water at 121 °C for 15 min. Then, 20 mL of 0.22 μm syringe-filtered 50% (w/v) glucose was added to the cooled media. 5-FOA media was prepared by autoclaving 250 mL of milli-Q water at 121 °C for 15 min. Yeast synthetic drop-out media without uracil was prepared with synthetic −Ura drop-out supplement from Sigma–Aldrich (0.96 g), yeast nitrogen base(without amino acids and with ammonia (3.35g), uracil (5 mg), 5-FOA in DMSO (5 mL of a 100 mg/mL solution), and glucose (10 g) in 250 mL of milli-Q water. The pH of this solution was adjusted to 4.0 with aqueous sodium hydroxide, and the resulting solution was filtered through a 0.22 μm filter. The filtered solution was added to the autoclaved water once it cooled to ~55 °C. To prepare solid media in 100 × 20 mm culture dishes from VWR, 20 g of bacteriological agar from VWR was added to each L of autoclaved solution. Approximately 10 mL of media was poured into each culture dish.

For liquid culture, *S. cerevisiae* cells were grown in 50 mL Falcon tubes from Corning Life Sciences (Tewksbury, MA, USA) at 30 °C with shaking at 200 rpm in 10 mL of YPD media. For culture on solid media, yeast cells were grown at 30 °C in an incubator until colonies were sufficiently large for replica plating. To keep the solid media from drying during long incubation times, a tray of sterilized milli-Q water was placed on the bottom rack of the incubator. Replica plating was performed using a replica plating tool for nutrient plates (SP Scienceware, Wayne, NJ, USA) and sterile velveteen squares (SP Scienceware). Alcohol burners were used to create a sterile benchtop work area during liquid and solid media preparation and the handling of yeast.

#### Transformation of *pdi1Δ Saccharomyces cerevisiae* and plasmid shuffling

We evaluated the ability of the CXC-containing plasmid to complement *pdi1*Δ *S. cerevisiae* using plasmid shuffling. The parent yeast strain used in this work, W303-1Bα Δ*pdi1*/pCT37, is a complete genetic knockout of wild-type *pdi1*. The cells of this yeast strain have been rescued with plasmid pCT37, which directs the expression of a non-genomically integrated *PDI* transgene, allowing for plasmid shuffling. Following the protocol of the Frozen-EZ Yeast Transformation Kit II from Zymo Research, the parent yeast was transformed with YEpWL.CXClinkerHDEL (*CXClinkerHDEL* and *TRP1*), YEpWL.RNaseA (*RNaseA* and *TRP1*; negative control), pRS424 (*S. cerevisiae PDI1* and *TRP1*; positive control), or no plasmid. To maintain a CXC library after transformation, yeast cells were grown overnight in liquid YPD media at 30 °C with shaking at 200 rpm. A 100 μL aliquot from the 10 mL overnight cultures was streaked onto YPD plates, which were then incubated at 30 °C for ~3 days until colonies appeared. All transformed yeast grew colonies on YPD plates.

After growth on YPD media, the transformed yeast was replica plated onto −Trp and −Ura plates. After growth, colonies from −Trp plates were replica plated onto 5-FOA and −Ura plates.

Lastly, after growth, colonies from 5-FOA plates were replica plated onto −Trp plates and −Ura plates. This cycle of replica plating was repeated four times. Successful complementation was assessed by the growth of a colony on 5-FOA and −Trp, but not on −Ura plates.

The largest *S. cerevisiae* +CXC-linker-HDEL −PDI colonies from successful plasmid shuffling plates were chosen for characterization (*i.e.*, doubling time, PCR, sequencing, microscopy, and RNA-seq). These colonies were streaked individually on 5-FOA and −Ura plates. Streaking verified complementation and ensured that individual clones were isolated from the initial yeast colonies. Three isolated colonies from each streaked plate were randomly selected as biological replicates. The triplicates were grown to mid-log phase in YPD media at 30 °C with shaking at 250 rpm and stored as glycerol stocks (0.22 μm syringe-filtered 30% (w/v) glycerol at −80 °C) for use in all in vivo characterization experiments.

**Polymerase chain reaction and agarose gel electrophoresis for genotyping**

Yeast strains were genotyped to determine whether PDI or a CXC-linker-HDEL peptide was present in complemented cells. Cultures of strains were grown to $1.0 \times 10^6$ cells in YPD media at 30 °C with shaking at 200 rpm, based on $OD_{600\ nm} = 1.0$ corresponding to $3 \times 10^7$ cells $mL^{-1}$ (*9*). Plasmid DNA was extracted following the protocol of the Zymoprep Yeast Plasmid Miniprep II kit from Zymo Research. The protocol of the KAPA2G Robust HotStart ReadyMix from Roche (Basel, Switzerland) was followed for PCR amplification of the genes of interest (table S8). The PCR product was directly loaded into a 1% w/v TAE Mini ReadyAgarose plus ethidium bromide Precast Gel from Bio-Rad. A 5 μL aliquot of DNA 1 kb Plus Ladder from New England Biolabs was added to one well. Gels were imaged on the ethidium bromide channel of a Chemidoc MP gel imaging station from Bio-Rad. Bands corresponding to the expected length of PDI or CXC-linker-HDEL fragments were cut out of the gel, and DNA was extracted from the gel following the protocol of the GeneJET Gel Extraction Kit from Thermo Fisher Scientific. Purified PCR products were sent to Quintara (Cambridge, MA, USA) for Sanger sequencing. The sequencing results were aligned with plasmid constructs using the online platform from Benchling (San Francisco, CA, USA).

**Doubling time**

Parent *S. cerevisiae* cells complemented with yeast PDI, CWC-α-helix-HDEL, and CRC-α-helix-HDEL were grown in YPD liquid medium. Three different clones from each cell type were analyzed. These cultures were diluted in YPD to a cell density of $OD_{600} = 0.10$ ($3 \times 10^6$ cells $mL^{-1}$). The resulting 10 mL cultures were grown in 50 mL Falcon tubes at 30 °C with shaking at 250 rpm. At 2-hour intervals, a 100 μL aliquot was removed and added to 900 μL of YPD media (1:10 dilution), and the $OD_{600}$ was measured. The doubling time of each culture was calculated as described previously (*1*). Doubling times were normalized to that of the parent yeast, which was 1.63 ± 0.08 hours.

**Microscopy and size determination**

Images were taken on a EVOS M7000 microscope using the Celleste 6 Image Analysis Software with a 60× 1.40 NA oil immersion objective from Thermo Fisher Scientific. 3,3′-Dihexyloxacarbocyanine iodide ($DiOC_6$;10 μM), a green fluorescent dye that localizes to phospholipid membranes (*10*), was added to cells 10 min before imaging. Images were taken in brightfield and green fluorescence channels (excitation and emission at 482 and 524 nm, respectively). Images were analyzed with FIJI ImageJ (*11*). Cell diameter was measured in triplicate and averaged for each sample.

**Thiol p$K_a$ of synthetic peptides**

Thiol p$K_a$ values for the CGC and CXC-linker-HDEL peptides were determined by measuring absorbance at 238 nm upon pH change at room temperature (*12*). Stock solutions of $KH_2PO_4$, $K_2HPO_4$, and $K_3PO_4$ (100 mM) were prepared to produce ten buffers with pH 5.5–12.0. These buffers were degassed with $N_2$(g), and the pH of each buffer was checked prior to experimentation. A 1.5 mM stock solution of a peptide in 100 mM $KH_2PO_4$ was prepared and degassed with $N_2$(g). Blank solutions for each pH were prepared by adding 70 µL of 100 mM $KH_2PO_4$ to 930 µL of each buffer. A 70 µL aliquot of the peptide stock solution was combined with 930 µL of each buffer, and the absorbance at 238 nm was measured at each pH in a quartz glass cuvette with a Cary 60 UV–vis spectrometer. The peptide mixture was recovered, and the pH was measured with an accumet XL500 pH meter from Thermo Fisher Scientific that was calibrated to pH 4.00, 7.00, and 10.00 on the day of use. The absorbance for each peptide was measured at each pH in triplicate. The mean values were fitted to a nonlinear sigmoidal variable slope curve using Prism v10.0.3 from GraphPad Software (Boston, MA, USA). Thiol p$K_a$ values were taken as the intersection of the second derivative curve with the abscissa (*13*), that is, the pH when $d^2$(fraction thiolate)/d(pH)$^2$ = 0 (fig. S9).

**Synthesis of disulfide bond isomerization assay substrate and product**

dns-sTI and dns-nTI were synthesized and purified as reported previously (*14*). After lyophilization, dns-sTI and dns-sTI were lyophilized was analyzed by Q-TOF ESI mass spectrometry (table S5).

The disulfide pairings in dns-nTI and dns-sTI were confirmed by proteolytic digestion with α-chymotrypsin. dns-nTI and dns-sTI (15 µg/mL) were incubated with α-chymotrypsin (0.5 µg/mL) for 18 hours at 37 °C in 100 mM Tris–HCl buffer, pH 7.6, containing $CaCl_2$ (1.0 mM). The digested samples were analyzed by mass spectrometry. As previously reported, α-chymotrypsin cleaves after the consecutive arginine residues in positions 14 and 15 in both peptides but is unable to cleave after phenylalanine in position 4 of sTI (*14*). After digestion, analysis by mass spectrometry yielded diagnostic fragments of dns-nTI (calculated 806.3 and 1148.9; found 807.4 and 1151.6) and dns-sTI (calculated 788.3 and 1148.5; found 789.4 and 1151.6 $[M + Cl]^-$).

**Disulfide bond isomerization assay with dns-sTI**

To minimize air oxidation during the assay, the reaction buffer (which was 100 mM Tris–HCl buffer, pH 7.6, containing 1.0 mM EDTA) was degassed under vacuum for 30 min and flushed with $N_2$(g) for 30 min. To reduce peptide adsorption to wells, 96-well black, flat-bottom polystyrene plates (Corning Life Sciences, Tewksbury, MA, USA) were prepared by soaking wells in 200 µL of 1% w/v aqueous poly(ethyleneimine) for at least 1 h, then rinsing with milli-Q water and allowing to air-dry prior to use.

CGC and CXC-α-helix-HDEL peptides (40 µM) were preincubated in reaction buffer at room temperature for 30 min. A solution (100 µL) of dns-sTI (2.2. µM) in the reaction buffer was added to wells in the plate. An aliquot (100 µL) of CGC or a CXC-α-helix-HDEL peptide was added to wells containing dns-sTI. Isomerization was measured over time by monitoring the increase in fluorescence at 465 nm upon excitation at 280 nm with a Spark multimode plate reader from Tecan (Männedorf, Switzerland). Reactions were run in triplicate.

Data from assays with dns-sTI alone and CGC or CXC-α-helix-HDEL peptides alone were averaged and used to background-correct data from assays with dns-sTI plus a peptide. The resulting data were used to calculate a second-order reaction rate constant, $k_2$, by fitting an asymptotic regression model using Prism. Values of $k_2$ were normalized to that for the CGC peptide, which had $k_2 = 84.8 \pm 1.0\ M^{-1}\ s^{-1}$. The $k_2$ values for the peptides are $10^1$- to $10^2$-fold lower than those for PDI or analogous enzymes (*14*, *15*).

**RNA-seq: Library preparation with poly(adenylic acid) selection and Illumina sequencing**
Aliquots (3 biological replicates) of *pdi1*Δ *S. cerevisiae* complemented with CRC-α-helix-HDEL, CWC-α-helix-HDEL, and the parent yeast strains, were sent to Azenta Life Sciences for RNA extraction and RNA sequencing. Total RNA was extracted from cells using Qiagen RNeasy Plus Universal Mini Kit following the manufacturer's instructions (Qiagen, Hilden, Germany). RNA samples were quantified using a Qubit 2.0 fluorometer from Thermo Fisher Scientific, and RNA integrity was confirmed with 4200 TapeStation from Agilent Technologies.

A strand-specific RNA sequencing library was prepared by following the protocol of a Next Ultra II Directional RNA Library Prep Kit from New England Biolabs. Briefly, the enriched RNA was fragmented for 8 min at 94 °C. Subsequently, first- and second-strand cDNA were synthesized. The second strand of cDNA was marked by incorporating dUTP during the synthesis. cDNA fragments were polyadenylated at the 3′ ends, and an index adapter was ligated to cDNA fragments. Limited-cycle PCR was used for library enrichment. The incorporated dUTP in the second-strand cDNA quenched amplification of the second strand, thereby preserving strand specificity. The sequencing library was validated on an Agilent TapeStation and quantified using a Qubit 2.0 fluorometer and quantitative PCR with reagents from KAPA Biosystems (Wilmington, MA, USA).

The sequencing libraries were multiplexed and clustered onto a flow cell on the Illumina NovaSeq instrument according to the manufacturer's instructions. The samples were sequenced using a 2 × 150 bp Paired End (PE) configuration.

Image analysis and base calling were conducted by the NovaSeq Control Software (NCS). Raw sequence data (.bcl files) generated from the Illumina NovaSeq were converted to fastq files and de-multiplexed using Illumina bcl2fastq 2.20. One mismatch was allowed for index sequence identification.

**RNA-seq: Data analysis**
After assessing the quality of the raw data, sequence reads were trimmed to remove possible adapter sequences and low-quality nucleotides using Trimmomatic v.0.36. The trimmed reads were mapped to the *S. cerevisiae* S288c reference genome available on ENSEMBL using STAR aligner v.2.5.2b, which uses a splice aligner that detects splice junctions and incorporates them to help align the entire read sequences. BAM files were generated from this step. Unique gene hit counts were calculated by using feature Counts from the Subread package v.1.5.2. Only unique reads that fell within exon regions were counted.

After extracting gene hit counts, the gene hit counts table was used for downstream differential expression analysis. Using DESeq2 (*16*), gene expression was compared between sample groups. The Wald test was used to generate *P*-values and $\log_2$-fold changes. Genes with adjusted *P*-values ($P_{adj}$) <0.05 and absolute $\log_2$-fold changes >1 were designated as differentially expressed genes for each comparison. A gene ontology analysis was performed on the statistically significant set of genes using the GeneSCF software. The mgi GO list was used to cluster the set of genes based on their biological process and determine their statistical significance. A PCA analysis was performed using the plotPCA function within the DESeq2 R package. The plot shows the samples in a 2D plane spanned by their first two principal components. The top 500 genes, selected by highest row variance, were used to generate the plot.

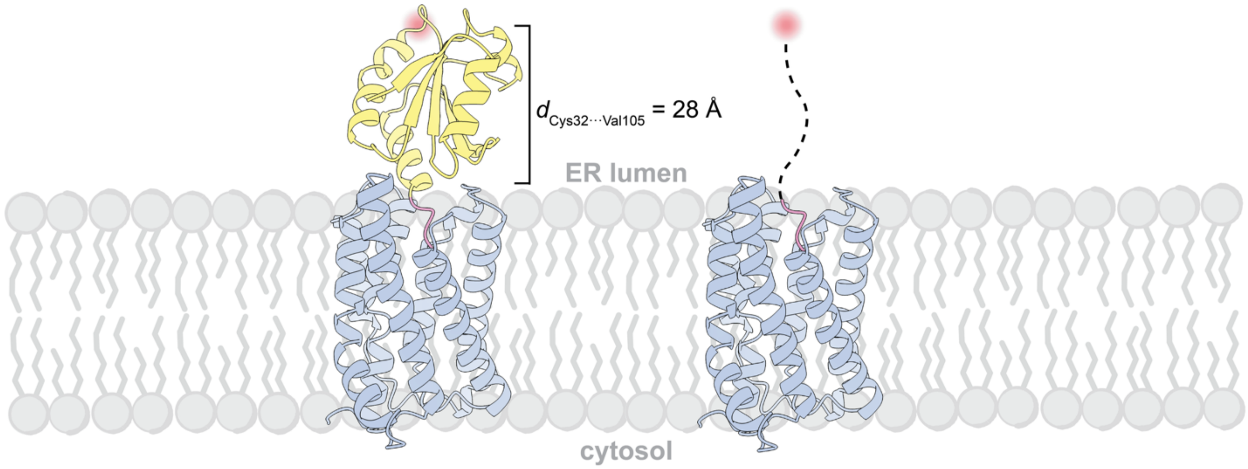


**Fig. S1. Estimation of the minimal linker length required for peptide-mediated replacement of PDI.** AlphaFold3-predicted structure of thioredoxin bearing a C-terminal HDEL sequence (yellow and pink; UniProt P10599) bound to the KDEL receptor (gray; UniProt P24390) in the endoplasmic reticulum membrane. The position of the N-terminal cysteine in the CGPC active site is highlighted in red. The distance between $C^{\alpha}$ of that residue (Cys32) and $C^{\alpha}$ of the thioredoxin residue N-terminal to the HDEL motif (Val105) is 28 Å. This distance was used to guide the design and length of CXC-linkers. The hypothetical linker is indicated by a dashed line.

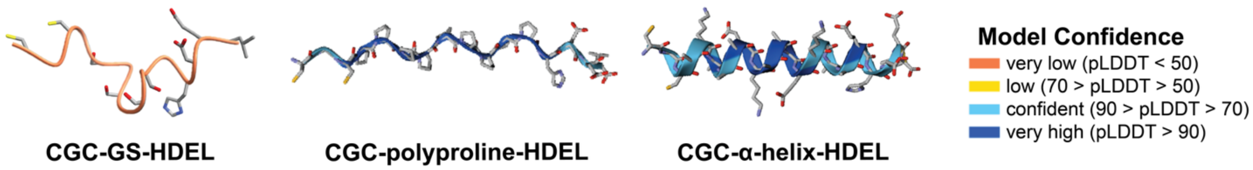


**Fig. S2. Predicted secondary structure of CXC-linker-HDEL peptides in solution.** AlphaFold3 predictions of CGC-GS-HDEL, CGC-polyproline-HDEL peptides, and CGC-α-helix-HDEL. Per-residue predicted local distance difference test (pLDDT) scores are shown, indicating low confidence for the flexible GS linker (pLDDT < 50) and higher confidence for the rigid polyproline and α-helix linkers (pLDDT > 70). Models were used to estimate linker geometry rather than to infer stable tertiary structure.

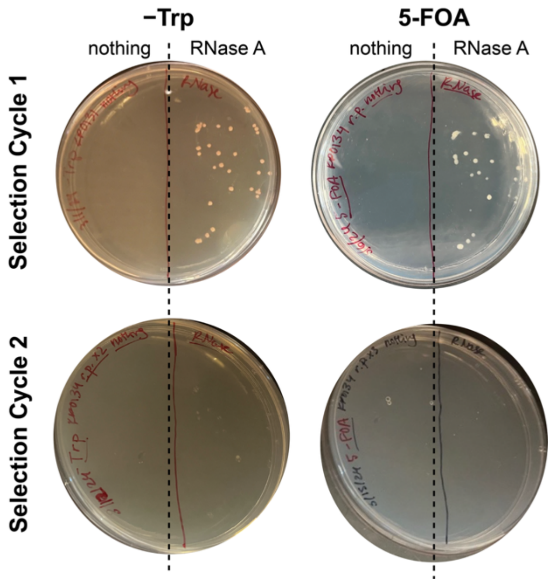


**Fig. S3. Negative controls for plasmid shuffling fail to replace PDI.** Plasmid shuffling of *pdi1*Δ *S. cerevisiae* transformed with either no second plasmid ("nothing") or a plasmid encoding bovine pancreatic ribonuclease A (RNase A). Replica plating from −Trp to 5-fluoroorotic acid (5-FOA) media shows loss of viability by the second selection cycle, confirming that unrelated proteins cannot replace PDI.

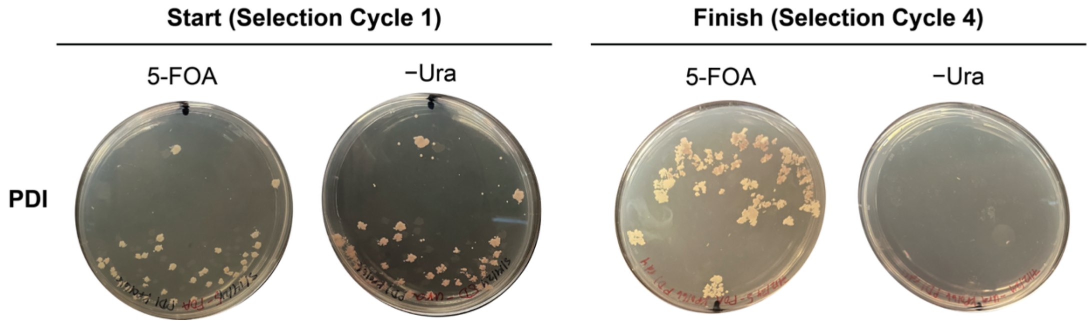


**Fig. S4. Positive control confirms plasmid shuffling strategy.** Plasmid shuffling of *pdi1*Δ *S. cerevisiae* transformed with a second plasmid encoding yeast PDI (*PDI1*, *TRP1*). Growth on 5-FOA and loss of growth on −Ura after four selection cycles confirms successful replacement of the original PDI plasmid.

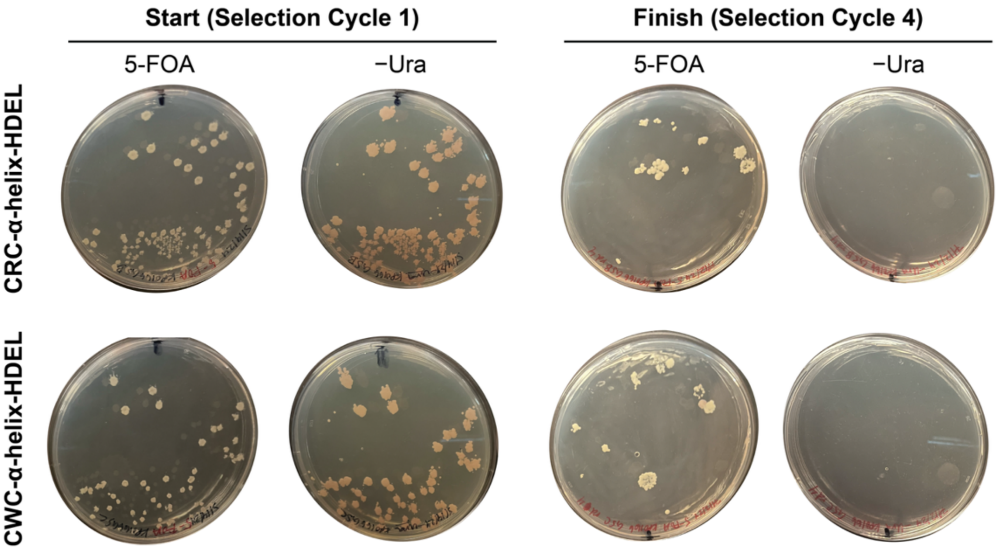


**Fig. S5. Identification of CXC and linker architectures capable of replacing PDI.** Plasmid shuffling results for CXC-GS-HDEL, CXC-polyproline-HDEL, and CXC-α-helix-HDEL peptide libraries. Viable *S. cerevisiae* colonies after four selection cycles were observed exclusively for CRC and CWC with α-helical linkers, indicating the requirements for peptide-mediated replacement of PDI.

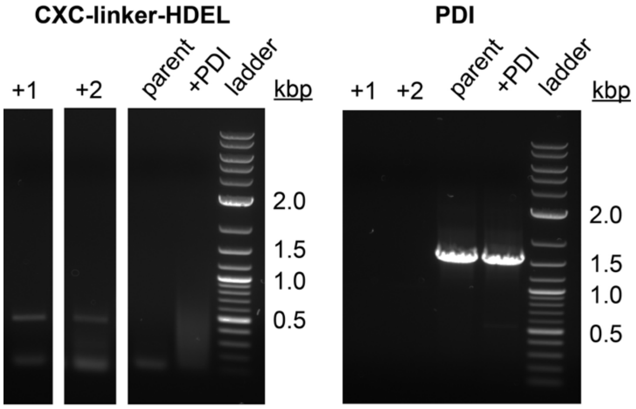


**Fig. S6. Genotyping confirms loss of PDI and retention of CXC-linker-HDEL constructs.** Polymerase chain reaction (PCR) analysis of genomic DNA from peptide-replaced yeast strains (+1 and +2), parent strain, and PDI-complemented control (+PDI). Amplification of CXC-linker-HDEL sequences (384 bp) and absence of PDI amplification (1566 bp) confirm replacement of PDI by peptide catalysts.

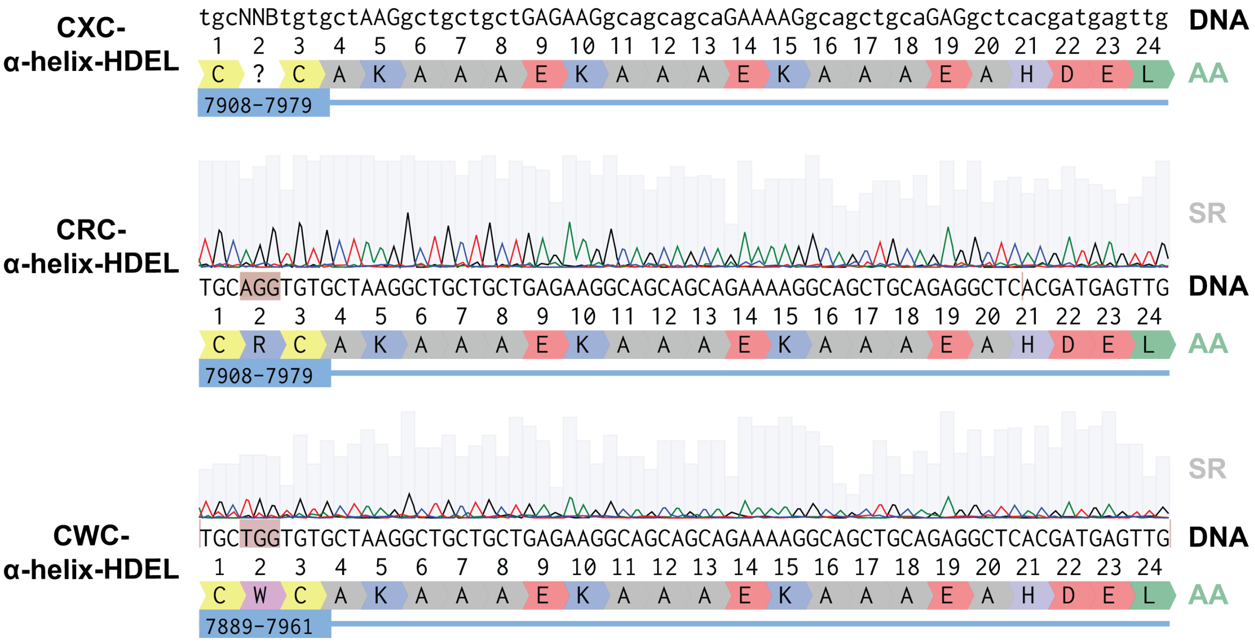


**Fig. S7. Sequence identification of peptides that replace PDI in vivo.** Sanger sequencing of PCR-amplified CXC-linker-HDEL constructs from peptide-replaced yeast. Alignments identify CRC-α-helix-HDEL and CWC-α-helix-HDEL as the peptides capable of replacing PDI. Quality scores (Q-scores) are shown for each nucleotide position.

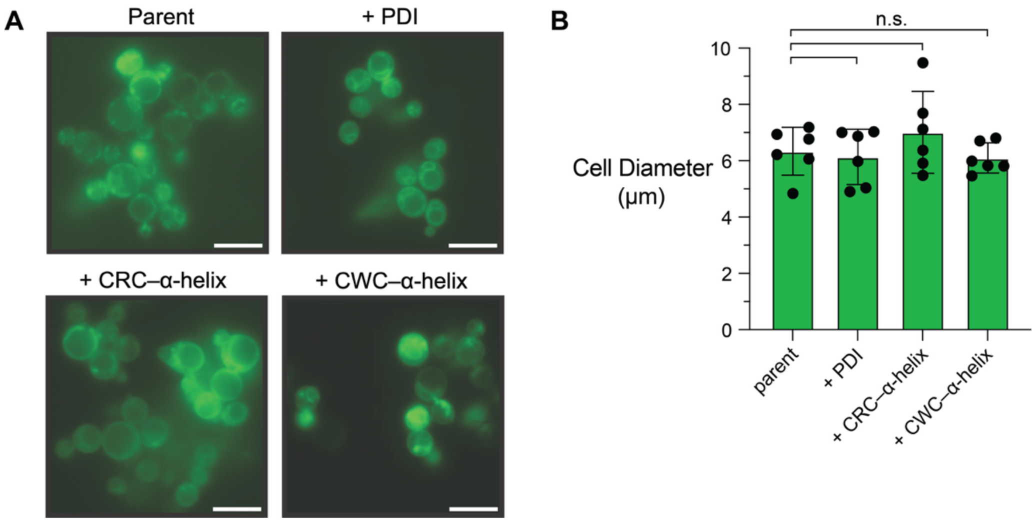


**Fig. S8. Morphology of peptide-replaced yeast cells.** (**A**) Fluorescence microscopy of parent, PDI-complemented, and peptide-replaced *S. cerevisiae* cells stained with 3,3′-dihexyloxacarbocyanine iodide ($DiOC_6$), which labels phospholipid membranes. Scale bar, 10 μm. (**B**) Quantification of cell diameter shows no significant difference between peptide-replaced and control strains ($P > 0.05$), indicating that replacement of PDI does not grossly alter cell morphology.

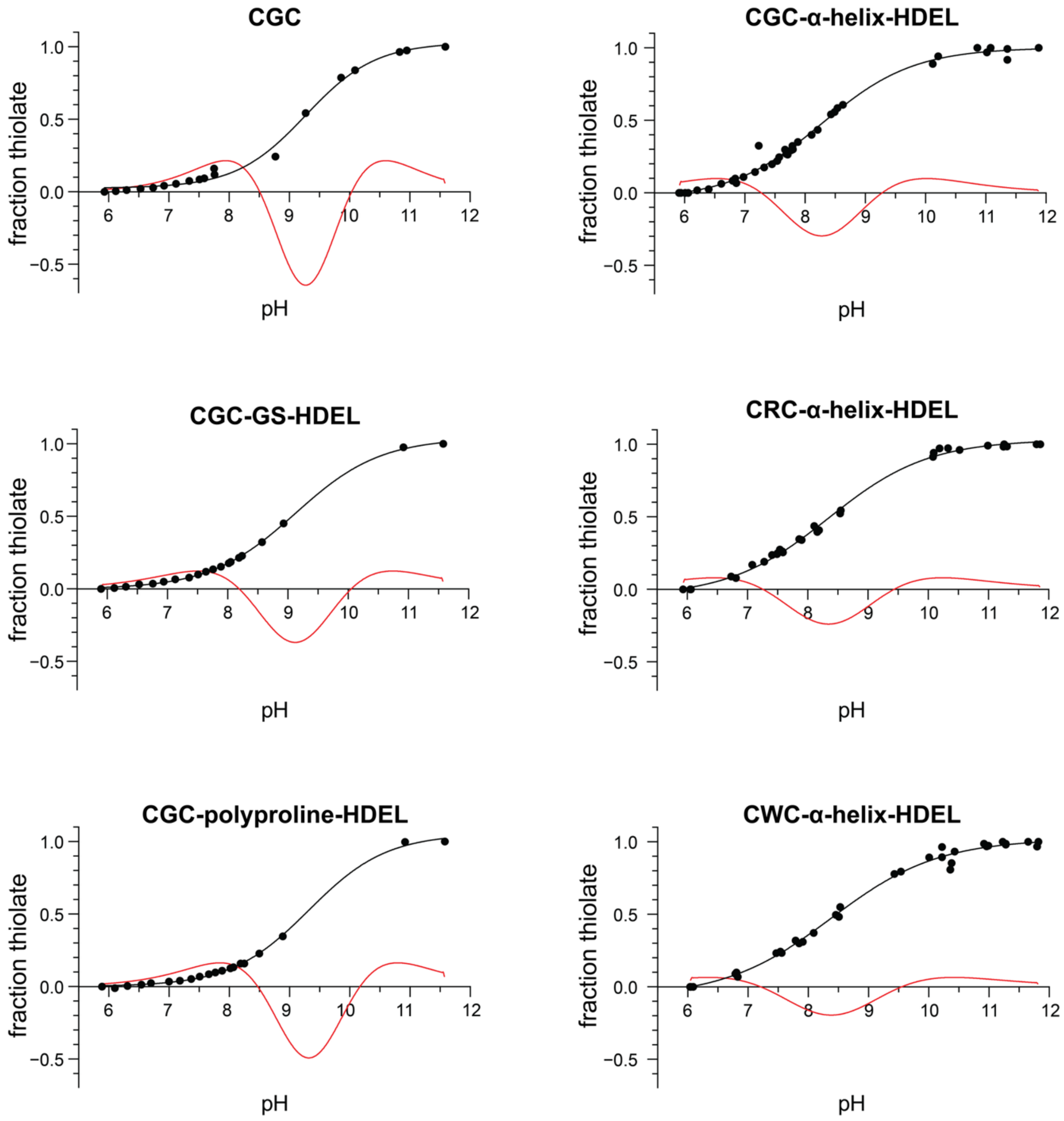


**Fig. S9. Thiol p$K_a$ values of CGC and CGC-linker-HDEL peptides.** Fraction thiolate was determined from the $A_{238\ nm}$ at different pH values (black). The p$K_a$ values for the two thiols in each peptide were the intersection of the second derivative curve (red) with the abscissa and are listed in Fig. 2B.

**Table S1. Amino acid sequences used for AlphaFold3-based estimation of linker geometry.** Amino acid sequences of CXC-linker-HDEL peptides, thioredoxin-HDEL, and the KDEL receptor were used to estimate linker length and orientation.

| Figure | Peptide or Protein | Amino Acid Sequence | UniProt ID |
|---|---|---|---|
| Fig. 1C | CGC-GS-HDEL | CGCGSGSGSGSGHDEL | — |
| | KDEL receptor | MNLFRFLGDLSHLLAIILLLLKIWKSRSCA<br>GISGKSQVLFAVVFTARYLDLFTNYISLYN<br>TCMKVVYIACSFTTVWLIYSKFKATYDGNH<br>DTFRVEFLVVPTAILAFLVNHDFTPLEILW<br>TFSIYLESVAILPQLFMVSKTGEAETITSH<br>YLFALGVYRTLYLFNWIWRYHFEGFFDLIA<br>IVAGLVQTVLYCDFFYLYITKVLKGKKLSL<br>PA | P24390 |
| Fig. 1C | CGC-polyproline-HDEL | CGCPPPPPPPPPHDEL | — |
| | KDEL receptor | MNLFRFLGDLSHLLAIILLLLKIWKSRSCA<br>GISGKSQVLFAVVFTARYLDLFTNYISLYN<br>TCMKVVYIACSFTTVWLIYSKFKATYDGNH<br>DTFRVEFLVVPTAILAFLVNHDFTPLEILW<br>TFSIYLESVAILPQLFMVSKTGEAETITSH<br>YLFALGVYRTLYLFNWIWRYHFEGFFDLIA<br>IVAGLVQTVLYCDFFYLYITKVLKGKKLSL<br>PA | P24390 |
| Fig. 1C | CGC-α-helix-HDEL | CGCAKAAAEKAAAEKAAAEAHDEL | — |
| | KDEL receptor | MNLFRFLGDLSHLLAIILLLLKIWKSRSCA<br>GISGKSQVLFAVVFTARYLDLFTNYISLYN<br>TCMKVVYIACSFTTVWLIYSKFKATYDGNH<br>DTFRVEFLVVPTAILAFLVNHDFTPLEILW<br>TFSIYLESVAILPQLFMVSKTGEAETITSH<br>YLFALGVYRTLYLFNWIWRYHFEGFFDLIA<br>IVAGLVQTVLYCDFFYLYITKVLKGKKLSL<br>PA | P24390 |
| Fig. S1 | thioredoxin | MVKQIESKTAFQEALDAAGDKLVVVDFSAT<br>WCGPCKMIKPFFHSLSEKYSNVIFLEVDVD<br>DCQDVASECEVKCMPTFQFFKKGQKVGEFS<br>GANKEKLEATINELVHDEL | P10599 |
| | KDEL receptor | MNLFRFLGDLSHLLAIILLLLKIWKSRSCA<br>GISGKSQVLFAVVFTARYLDLFTNYISLYN<br>TCMKVVYIACSFTTVWLIYSKFKATYDGNH<br>DTFRVEFLVVPTAILAFLVNHDFTPLEILW<br>TFSIYLESVAILPQLFMVSKTGEAETITSH<br>YLFALGVYRTLYLFNWIWRYHFEGFFDLIA<br>IVAGLVQTVLYCDFFYLYITKVLKGKKLSL<br>PA | P24390 |

**Table S2. Estimated linker lengths for CXC-linker-HDEL peptides.** Linker lengths were estimated based on average residue length for flexible linkers and average rise per residue for polyproline II and α-helix conformations. Estimates were used to guide construct design for peptide-mediated replacement of PDI.

| Peptide | Amino Acid Sequence | Average Length of a Residue* or Average Rise per Helical Residue† | Number of Residues in CXC-linker | Estimated CXC-linker Length |
|---|---|---|---|---|
| CGC-GS-HDEL | CGCGSGSGHDEL | 3.15 Å* | 8 | 25 Å |
| CGC-polyproline-HDEL | CGCPPPPPPPPPPHDEL | 3.10 Å† | 12 | 37 Å |
| CGC-α-helix-HDEL | CGCAKAAAEKAAAEKAAAEAHDEL | 1.50 Å† | 20 | 30 Å |

*Average length per residue assumes an extended conformation.
†Average rise per helical residue corresponds to an idealized polyproline II or α-helix geometry.

**Table S3. Amino acid sequences of synthetic peptides used in biochemical and cellular assays.** Sequences include CGC and CXC-linker-HDEL constructs tested for catalytic activity and replacement of PDI in vivo.

| Peptide | Amino Acid Sequence |
|---|---|
| CGC | H-**CGC**-$NH_2$ |
| CGC-GS-HDEL | H-**CGCGSGSGHDEL**-OH |
| CGC-polyproline-HDEL | H-**CGCPPPPPPPPPHDEL**-OH |
| CGC-α-helix-HDEL | H-**CGCAKAAAEKAAAEKAAAEAHDEL**-OH |
| CRC-α-helix-HDEL | H-**CRCAKAAAEKAAAEKAAAEAHDEL**-OH |
| CWC-α-helix-HDEL | H-**CWCAKAAAEKAAAEKAAAEAHDEL**-OH |
| dns-sTI | Ac-**KWCFRVCYRGICYRRCK(dns)G**-OH |
| dns-nTI | Ac-**KWCFRVCYRGICYRRCK(dns)G**-OH |

**Table S4. High-performance liquid chromatography (HPLC) methods used for purification of synthetic peptides.** Peptides were purified using an XSelect Peptide C18 preparatory column in Phase A/Phase B gradients at ambient temperature. Phase A: 95% $H_2O$/5% ACN/0.1% TFA; Phase B: 95% CAN/5% $H_2O$/0.1% TFA.

| **HPLC Method 1 (for CGC)** | | | |
|---|---|---|---|
| **Time (min)** | **A (%)** | **B (%)** | **Flow (mL/min)** |
| 0 | 100.00 | 0.00 | 6.00 (4.00 for CRC) |
| 4 | 100.00 | 0.00 | 6.00 (4.00 for CRC) |
| 10 | 0.00 | 100.00 | 6.00 (4.00 for CRC) |
| 12 | 100.00 | 0.00 | 6.00 (4.00 for CRC) |
| 15 | 100.00 | 0.00 | 6.00 (4.00 for CRC) |
| **HPLC Method 2 (for CXC-linker-HDEL peptides)** | | | |
| **Time (min)** | **A (%)** | **B (%)** | **Flow (mL/min)** |
| 0 | 100.00 | 0.00 | 14.00 |
| 4 | 100.00 | 0.00 | 14.00 |
| 10 | 0.00 | 100.00 | 14.00 |
| 12 | 0.00 | 100.00 | 14.00 |
| 15 | 100.00 | 0.00 | 14.00 |
| **HPLC Method 3 (for dns-sTI and dns-nTI)** | | | |
| **Time (min)** | **A (%)** | **B (%)** | **Flow (mL/min)** |
| 0 | 72.00 | 28.00 | 14.60 |
| 30 | 39.00 | 61.00 | 14.60 |
| 32 | 0.00 | 100.00 | 14.60 |
| 34 | 0.00 | 100.00 | 14.60 |
| 35 | 72.00 | 28.00 | 14.60 |

**Table S5. Mass spectrometry validation of synthetic peptides.** Calculated and observed *m/z* values and retention times confirm the identity and purity of peptides used in biochemical and cellular experiments.

| Peptide | *m/z* | | HPLC | |
|---|---|---|---|---|
| | Calculated | Observed | Retention Time (min) | Method* |
| CGC | 280.06 | 281.07 | 3.1 | 1 |
| CGC-GS-HDEL | 1120.38 | 1121.40 | 9.2 | 2 |
| CGC-polyproline-HDEL | 1648.74 | 1650.75 | 9.0 | 2 |
| CGC-α-helix-HDEL | 2328.07 | 2330.09 | 9.2 | 2 |
| CRC-α-helix-HDEL | 2428.15 | 2430.18 | 9.1 | 2 |
| CWC-α-helix-HDEL | 2457.13 | 2460.16 | 9.2 | 2 |
| dns-nTI | 2497.1 | 2502.6 | 9.3 | 3 |
| dns-sTI | 2491.1 | 2499.9 | 9.3 | 3 |

*HPLC method refers to table S4.

**Table S6. DNA sequences used to construct CXC-linker-HDEL peptide libraries.** Degenerate codons (NNB) were used to encode all 20 canonical amino acids at the X position while minimizing stop codons. N = a, c, g, or t; B = c, g, or t.

| Peptide Library | DNA Sequence |
|---|---|
| CXC-GS-HDEL | tgcNNBtgtggttccggctctggttctggttctggtcacgatgagttgtaa |
| CXC-polyproline-HDEL | tgcNNBtgtccacctccgccacctccgccacctccgcacgatgagttgtaa |
| CXC-α-helix-HDEL | tgcNNBtgtgctaaggctgctgctgagaaggcagcagcagaaaaggcagctgcaga ggctcacgatgagttgtaa |

**Table S7. Oligonucleotide primers used for genotyping of yeast strains with the polymerase chain reaction (PCR).**

| Encoded PCR Product | Primer | Primer Sequence (5′→3′) | PCR Product Length |
|---|---|---|---|
| **CXC-linker-HDEL** | Forward | cacacataaacaaacaccatggga | 384 bp |
| | Reverse | gtcgacggtatcgataagct | |
| **PDI** | Forward | atgaagttttctgctggtgc | 1566 bp |
| | Reverse | caattcatcgtgaatggcatcttc | |

**Table S8. Thermocycler conditions used for PCR amplification with the polymerase chain reaction (PCR).**

| PCR of CXC-linker-HDEL | | |
|---|---|---|
| *T* (°C) | Time (min:s) | Cycles |
| 95 | 3:00 | |
| 95 | 0:15 | Cycle 45× |
| 60 | 0:15 | |
| 72 | 0:15 | |
| 72 | 2:00 | |
| 4 | infinite hold | |
| **PCR of PDI** | | |
| *T* (°C) | Time (min:s) | Cycles |
| 95 | 1:00 | |
| 95 | 0:15 | Cycle 35× |
| 60 | 0:15 | |
| 72 | 0:15 | |
| 72 | 5:00 | |
| 4 | infinite hold | |

**Table S9. Differentially expressed genes associated with translational processes following replacement of PDI.** $Log_2$-fold changes in expression of genes associated with ribosome biogenesis, cytosolic translation, and mitochondrial translation in peptide-replaced yeast relative to the parent strain. Differential expression is defined as $P_{adj} < 0.05$.

| Ribosome biogenesis | | | Cytosolic translation | | | Mitochondrial translation | | |
|---|---|---|---|---|---|---|---|---|
| gene | CWC log2foldchange | CRC log2foldchange | gene | CWC log2foldchange | CRC log2foldchange | gene | CWC log2foldchange | CRC log2foldchange |
| ACL4 | -1.197563516 | -0.647561643 | AFG3 | 1.21640437 | 0.777458465 | 21S_RRNA | -9.115736819 | -11.12082554 |
| ALB1 | -2.00343816 | 1.510629886 | LHS1 | 1.434362376 | 1.544396362 | 15S_RRNA | -6.519748407 | -11.48394654 |
| ARB1 | 1.00852042 | 0.860857591 | MRPL10 | 2.660749221 | 1.408358858 | IFM1 | 2.500187209 | 1.464947746 |
| BCD1 | -1.636079308 | -1.474471528 | MRPL25 | 1.874649584 | 1.705245259 | IMG1 | 3.019353726 | 1.473977606 |
| BRX1 | -2.155806684 | 0.638145672 | MRPL28 | 2.124058606 | 1.397247998 | IMG2 | 1.91460477 | 1.426428079 |
| BUD21 | -1.746237983 | -0.598759307 | MRPL32 | 2.52688499 | 2.4528037 | ISM1 | 1.82662887 | 1.363283938 |
| CIC1 | -1.864740934 | 0.83508701 | MRPL38 | 0.933503929 | 1.006877192 | MEF1 | 3.597641741 | 3.126503603 |
| DBP2 | -3.538124095 | 0.523781189 | MRPL6 | 1.683003886 | 1.805862344 | MEF2 | 2.956273964 | -1.679441593 |
| DBP6 | -1.582027719 | -0.092448556 | MRPS35 | 2.840864909 | 2.716374983 | MHR1 | 1.85461132 | 1.683992362 |
| DBP7 | -2.313166535 | 0.369826721 | MRPS5 | 1.705980649 | 1.227004434 | MNP1 | 2.744798012 | 2.616317274 |
| DBP9 | -1.233784449 | 0.775477957 | MTG2 | 3.835093716 | 1.642533934 | MRF1 | 1.468608914 | 1.90060742 |
| DHR2 | -1.124160664 | 1.539988158 | RBG2 | -2.100535346 | 0.167795038 | MRP1 | 3.14850107 | 2.333469817 |
| DIP2 | -1.871510882 | 1.099178125 | RML2 | 3.101696642 | 2.326840173 | MRP10 | 1.237146683 | 1.135354626 |
| ECM1 | -2.059358082 | 0.145814415 | RPL10 | -2.110267842 | -0.490310893 | MRP13 | 3.639728536 | 3.334122617 |
| ECM10 | -0.955671986 | -1.537668069 | RPL12A | -1.335164417 | 1.278119963 | MRP17 | 2.343825825 | 1.807031548 |
| ECM11 | -1.723848071 | -1.306345587 | RPL12B | -1.757968448 | 1.033630779 | MRP2 | 2.352032315 | 2.328859579 |
| ECM12 | -1.589590719 | -0.699451895 | RPL13A | -1.779815277 | 0.523105175 | MRP20 | 1.884523246 | 0.785765768 |
| ECM13 | -1.062650151 | -1.442159351 | RPL13B | -1.263701525 | 0.799259943 | MRP21 | 1.217105593 | 1.278662923 |
| ECM14 | 0.564946826 | -0.296518536 | RPL14A | -1.821687668 | 1.481146281 | MRP4 | 1.538199345 | 1.372612494 |
| ECM15 | -1.269314166 | -1.07698444 | RPL14B | -1.225956846 | 0.415260607 | MRP49 | 1.371071114 | 1.397073159 |
| ECM16 | -0.773572898 | 0.595092522 | RPL16A | -2.526936922 | 0.501694374 | MRP51 | 2.055990874 | 2.143716864 |
| ECM18 | 0.654772844 | 0.027541791 | RPL16B | -1.31920782 | 1.269647886 | MRP7 | 3.113492483 | 1.853877477 |
| ECM19 | 0.24341868 | 0.577648672 | RPL17A | -1.304815946 | 1.298349985 | MRPL10 | 2.660749221 | 1.408358858 |
| ENP2 | -1.856922062 | 0.160290265 | RPL17B | -1.784017343 | 0.268125629 | MRPL11 | 2.687712737 | 2.589263208 |
| ERB1 | -1.520730663 | 0.993695156 | RPL18A | -1.624128539 | 1.02920594 | MRPL13 | 2.140649572 | 1.805384679 |
| ESF2 | -2.207693047 | -0.000884163 | RPL18B | -2.190367753 | -0.107392547 | MRPL15 | 2.89406995 | 2.591554819 |
| FAL1 | -1.095890048 | 0.66644199 | RPL19A | -1.425961991 | 0.78245857 | MRPL16 | 1.819568126 | 1.133802817 |
| FCF1 | -2.377053566 | -0.886771744 | RPL19B | -1.817561056 | 0.849932579 | MRPL17 | 2.068040121 | 1.855189174 |
| HAS1 | -1.508541751 | 1.117407161 | RPL20A | -1.747269413 | 1.021428201 | MRPL19 | 1.601740959 | 1.280754031 |
| HCA4 | -1.818525331 | 0.47868731 | RPL20B | -1.392300025 | 0.471656344 | MRPL20 | 3.200485765 | 2.689818514 |
| IMP3 | -1.639030105 | 0.820358707 | RPL21A | -2.087991259 | 1.13315671 | MRPL22 | 1.840393237 | 1.55982031 |
| IPI1 | -1.218145669 | 1.19506159 | RPL22A | -1.428437664 | 2.165982249 | MRPL23 | 1.926655114 | 1.729974735 |
| KRR1 | -2.866197809 | -0.684400006 | RPL23A | -1.646945455 | 0.80103215 | MRPL24 | 2.249833232 | 1.718656352 |
| MAK11 | -1.554957362 | -1.234536234 | RPL23B | -1.598855703 | 0.922364713 | MRPL27 | 1.635896545 | 1.067508369 |
| MAK21 | -1.440161423 | 0.934616652 | RPL24A | -2.282087934 | 0.527333537 | MRPL28 | 2.124058606 | 1.397247998 |
| MRPL11 | 2.687712737 | 2.589263208 | RPL24B | -1.139378142 | 1.162872847 | MRPL3 | 2.755372434 | 2.962758854 |
| NAN1 | -1.423094994 | 1.042054486 | RPL25 | -2.178724972 | 0.723950061 | MRPL31 | 2.95520166 | 2.148708319 |
| NIP7 | -2.459668736 | 1.205657927 | RPL26A | -1.628797012 | 0.830861698 | MRPL32 | 2.52688499 | 2.4528037 |
| NOC3 | -1.281502323 | 0.506313114 | RPL26B | -1.57952349 | 1.242562832 | MRPL33 | 1.602607628 | 0.77634018 |
| NOC4 | -1.720185963 | 0.883029259 | RPL27B | -2.747639062 | 0.700924324 | MRPL35 | 2.526915289 | 2.369414225 |
| NOG1 | -1.908608865 | 0.548263754 | RPL28 | -1.462127937 | 1.644276655 | MRPL36 | 2.417183956 | 2.148863461 |
| NOG2 | -1.797901981 | 1.278283039 | RPL29 | -1.039484711 | 1.571329944 | MRPL37 | 1.974235344 | 1.61429417 |
| NOP1 | -1.518025197 | 2.054487807 | RPL2A | -1.913119523 | 1.117112696 | MRPL38 | 0.933503929 | 1.006877192 |
| NOP10 | -2.184458556 | -0.453576594 | RPL31A | -1.723476557 | 1.303150134 | MRPL39 | 2.457868364 | 2.171749747 |
| NOP12 | -1.406860639 | 0.686207314 | RPL31B | -2.955724836 | 0.27459196 | MRPL4 | 2.674591328 | 2.538084547 |
| NOP13 | -0.760697767 | 1.365110126 | RPL32 | -1.628959947 | 1.118364577 | MRPL40 | 2.158567735 | 1.779491579 |
| NOP14 | -1.807560012 | 0.312470132 | RPL33A | -1.60571608 | 1.497670344 | MRPL44 | 0.794464023 | 0.930234065 |
| NOP15 | -3.850164973 | -0.603269385 | RPL33B | -2.567632419 | 0.151303622 | MRPL49 | 3.248378683 | 2.788129898 |
| NOP16 | -1.033399185 | -1.623148107 | RPL34A | -1.205669646 | 0.673169706 | MRPL50 | 2.303370289 | 2.416138707 |
| NOP19 | -2.128023899 | -0.740270682 | RPL36A | -1.82080694 | 1.019855828 | MRPL51 | 1.095911505 | 0.592543974 |
| NOP8 | -1.65586248 | 0.20376576 | RPL37A | -2.126069536 | 0.865660964 | MRPL6 | 1.683003886 | 1.805862344 |
| NOP9 | -1.779032809 | 0.35254597 | RPL38 | -1.273591723 | 1.12002516 | MRPL7 | 3.119625598 | 2.711017835 |
| NPL3 | 2.42213544 | -0.81425661 | RPL40A | -1.678075924 | 0.305465287 | MRPL8 | 2.990544992 | 1.658349619 |
| NSA2 | -3.691202478 | -0.107746571 | RPL40B | -3.028829589 | 0.019822113 | MRPL9 | 2.895711014 | 1.838099664 |
| NUG1 | -2.040244923 | 0.50104415 | RPL41B | -2.119704625 | 0.93669465 | MRPS12 | 2.105825815 | 1.303266338 |
| PWP2 | -2.363131891 | 0.278135527 | RPL42A | -1.979131814 | 1.170262747 | MRPS17 | 3.060648215 | 2.532524179 |
| RIA1 | -1.459314048 | -0.026250779 | RPL42B | -1.492081951 | 1.182791482 | MRPS18 | 1.598237706 | 0.875961622 |
| RIX1 | -1.427801728 | 0.642097978 | RPL43B | -1.646755775 | 2.175221087 | MRPS28 | 1.734160976 | 1.15863456 |
| RIX7 | -1.93416828 | 1.010821654 | RPL6A | -1.952419125 | 0.8674034 | MRPS35 | 2.840864909 | 2.716374983 |
| RLP24 | -1.39770163 | 0.900268526 | RPL6B | -1.260026229 | 2.428744527 | MRPS5 | 1.705980649 | 1.227004434 |
| ROK1 | -1.303687799 | 0.163130399 | RPL7A | -1.467833308 | 0.834983047 | MRPS8 | 2.666986461 | 2.026799755 |
| RPA12 | -1.603913322 | 1.065556829 | RPL7B | -1.558565065 | 0.342916769 | MRPS9 | 2.759989201 | 2.858355204 |
| RPA43 | -3.062475829 | 0.575790944 | RPL8A | -1.883573186 | 1.011132164 | MRX14 | 2.278580763 | 1.931971786 |
| RPA49 | -2.029350656 | 1.039653674 | RPL8B | -1.332709102 | 1.32533906 | MSE1 | 2.006387033 | 1.966423546 |
| RPC19 | -1.232980397 | 0.263525239 | RPM2 | -1.42277809 | -1.906705927 | MSK1 | 3.24854664 | 2.686404817 |
| RPC40 | -1.996261938 | 0.652423454 | RPP1A | -1.536337135 | 1.574288269 | MSR1 | 1.376367726 | -0.016506298 |
| RPL34A | -1.205669646 | 0.673169706 | RPP1B | -1.62833274 | 1.017787837 | NAM2 | 2.98467348 | 2.103779406 |
| RPL40A | -1.678075924 | 0.305465287 | RPS0A | -2.240477061 | 0.026887537 | NAM9 | 3.277557189 | 3.287761233 |
| RPL40B | -3.028829589 | 0.019822113 | RPS0B | -2.061799587 | 0.989865943 | PET123 | 2.657769806 | 2.928867244 |
| RPL8A | -1.883573186 | 1.011132164 | RPS11A | -2.55286705 | 0.093007849 | PPE1 | -1.284181816 | -1.522877522 |
| RPL8B | -1.332709102 | 1.32533906 | RPS13 | -1.588380759 | 1.651463275 | PTH1 | 2.939719246 | 0.311336082 |
| RPO26 | -2.003818259 | 0.294727061 | RPS14B | -1.949677442 | 0.770436982 | RML2 | 3.101696642 | 2.326840173 |
| RPS0A | -2.240477061 | 0.026887537 | RPS15 | -1.186908419 | 1.160028749 | RSM18 | 2.283853491 | 1.099202347 |
| RPS0B | -2.061799587 | 0.989865943 | RPS16A | -1.301691103 | 0.698219571 | RSM19 | 1.95673606 | 1.180336632 |
| RPS14B | -1.949677442 | 0.770436982 | RPS16B | -1.806592473 | 1.188348938 | RSM23 | 2.159750305 | 1.704545462 |
| RPS15 | -1.186908419 | 1.160028749 | RPS17B | -1.602607862 | 1.189430175 | RSM24 | 1.656798616 | 1.459480411 |
| RPS19A | -1.513580732 | 1.200253763 | RPS18B | -2.373695175 | 0.744554556 | RSM25 | 2.589301233 | 2.224331715 |
| RPS6A | -1.511208627 | 0.644875015 | RPS19A | -1.513580732 | 1.200253763 | RSM26 | 3.043970265 | 2.920518258 |
| RPS6B | -1.488672577 | 0.456942824 | RPS1A | -2.45769945 | -0.083418075 | RSM27 | 1.46043381 | 1.113699614 |
| RPS7A | -1.834445349 | 0.909618704 | RPS1B | -1.977233409 | 0.861229329 | RSM28 | 3.088898187 | 2.670138133 |
| RPS7B | -1.504931453 | 1.019421967 | RPS21B | -1.214150187 | 1.360855918 | RSM7 | 1.830180834 | 0.883965991 |
| RPS9B | -1.498147708 | 1.746261557 | RPS22A | -1.365103498 | 2.187807163 | RTC6 | 1.252868464 | 0.675674652 |
| RRP12 | -1.35535516 | 0.647849168 | RPS24A | -1.685338593 | 0.655009877 | SWS2 | 1.619538143 | 1.316651694 |
| RRP3 | -1.64452167 | 0.525162227 | RPS24B | -2.075691095 | 0.703257058 | TUF1 | 2.222258983 | 1.849572253 |
| RRP36 | -2.069274117 | -0.497358373 | RPS25A | -1.435899935 | 0.75649802 | YML6 | 1.741404077 | 1.749812355 |
| RRP5 | -1.01506443 | 0.678157061 | RPS25B | -1.9464518 | 0.399765807 | YMR31 | -1.333963881 | -1.659790536 |
| RRS1 | -2.209778078 | -0.439251631 | RPS26A | -1.384507451 | 1.452833026 | YMR310C | -0.94700368 | 0.257836916 |
| RTC6 | 1.252868464 | 0.675674652 | RPS26B | -1.35863579 | 1.937071432 | YMR315W | -0.760870573 | 0.476988009 |
| SLX9 | -1.747531805 | 0.441070263 | RPS27B | -1.710054091 | 0.395732522 | YMR315W-A | 0.543213785 | 1.983137143 |
| SNU13 | 1.393484964 | 3.626064609 | RPS29A | -1.584746481 | 0.816321812 | YMR316C-A | -0.243964158 | 1.727070141 |
| SSF2 | 1.102420796 | 0.53279331 | RPS29B | -1.906913388 | 0.431814563 | YMR317W | 0.419344027 | -1.15240021 |
| TMA20 | -1.89125584 | -0.60963408 | RPS3 | -1.183090075 | 1.454459651 | | | |
| TOR2 | -1.295319335 | -0.267979099 | RPS30A | -0.19437084 | 0.556153283 | | | |
| TSR1 | -2.264632823 | -0.003029531 | RPS30B | -1.647577377 | 0.258380727 | | | |
| TSR3 | -1.848121765 | -0.673467238 | RPS31 | -0.690525675 | 1.956764416 | | | |
| URB1 | -1.462298008 | 1.162387879 | RPS4A | -1.943752124 | 0.953406992 | | | |
| UTP10 | -1.482297754 | 1.37214471 | RPS4B | -2.313299538 | -0.068222382 | | | |
| UTP11 | -1.658980222 | -0.548904216 | RPS5 | -2.026768206 | 0.430869604 | | | |
| UTP13 | -2.079865556 | 1.551815378 | RPS6A | -1.511208627 | 0.644875015 | | | |
| UTP14 | -1.643140644 | 0.729731938 | RPS6B | -1.488672577 | 0.456942824 | | | |
| UTP18 | -1.487588732 | 1.162090731 | RPS7A | -1.834445349 | 0.909618704 | | | |
| UTP20 | -2.078797033 | 0.999929588 | RPS7B | -1.504931453 | 1.019421967 | | | |
| UTP21 | -1.468190211 | 1.347078307 | RPS8A | -1.57734924 | 0.568508261 | | | |
| UTP25 | -1.136399919 | 0.864988219 | RPS8B | -1.542330372 | 0.763998663 | | | |
| UTP4 | -2.014806913 | 0.718055961 | SLH1 | -1.075717501 | 0.504905846 | | | |
| UTP6 | -1.345365257 | 0.759804642 | SSB2 | -1.538653965 | 0.181923297 | | | |
| YTM1 | -1.387400483 | 0.829805988 | SUP19 | 1.399642568 | -1.028662441 | | | |
| | | | TIF3 | -1.906147276 | -0.990219089 | | | |
| | | | TIF34 | -0.120733144 | 0.319005419 | | | |
| | | | TIF35 | 0.059678122 | -0.008389114 | | | |
| | | | TMA7 | -1.236660895 | -0.059567323 | | | |

**Table S10. Gene ontology (GO) analysis of biological processes altered following PDI replacement.** GO categories enriched among differentially expressed genes in peptide-replaced yeast compared to the parent strain.

| Process | CWC-α-helix-HDEL | | CRC-α-helix-HDEL | |
|---|---|---|---|---|
| | ***P*-value** | **$P_{adj}$-value** | ***P*-value** | **$P_{adj}$-value** |
| **ribosome biogenesis** | 4.55E−05 | 0.21714096 | 0.99867593 | 0.999228691 |
| **cytoplasmic translation** | 4.48E−08 | 2.85E−05 | 0.99662181 | 0.999228691 |
| **mitochondrial translation** | 5.36E−09 | 5.11E−06 | 0.01044348 | 0.70325182 |

**Table S11. Differential expression of unfolded protein response and chaperone-related genes.** $Log_2$-fold changes and significance values for genes associated with the unfolded protein response (UPR; red), DnaJ-family chaperones (purple), and ER-associated degradation pathways (blue) in peptide-replaced yeast. n.d.: not determined.

| Gene | CWC-α-helix-HDEL | | | CRC-α-helix-HDEL | | |
|---|---|---|---|---|---|---|
| | $log_2$-fold change | *P*-value | $P_{adj}$-value | $log_2$-fold change | *P*-value | $P_{adj}$-value |
| **HAC1** | 1.16066236 | 0.00029686 | 0.00276492 | 2.70195757 | 0.00013502 | 0.00278745 |
| **KAR2** | 1.91942522 | 4.1206E−05 | 0.00029686 | 1.80223893 | 0.0012365 | 0.01473423 |
| **ERO1** | 1.75939699 | 0.00019464 | 0.00111063 | 2.36323119 | 9.0572E−06 | 0.00030196 |
| **LHS1** | 1.43436238 | 0.00598857 | 0.01961639 | 1.54439636 | 0.01679079 | 0.09320899 |
| **EUG1** | 0.59650893 | 0.11565353 | 0.21163999 | 0.62534276 | 0.04143494 | 0.16846998 |
| **INO1** | 0.7142104 | 0.11278575 | 0.20720864 | 0.01071204 | 0.98117659 | 0.99293482 |
| **JEM1** | 0.20720864 | 1.4540E−07 | 2.1729E−06 | 1.02778564 | 0.03040102 | 0.13796327 |
| **SCJ1** | 1.18135866 | 0.01364655 | 0.03840756 | 2.1722858 | 0.01316952 | 0.07991252 |
| **MDJ1** | 2.71150776 | 5.5959E−11 | 1.9989E−09 | 1.82157807 | n.d. | n.d. |
| **YDJ1** | 0.25262063 | 0.54821323 | 0.67638734 | 0.56600849 | 0.25815014 | 0.50214681 |
| **DER1** | −0.1184721 | 0.80173325 | 0.87114176 | 0.50214681 | 0.24350921 | 0.48543554 |
| **HRD1** | 0.59092423 | 0.15377542 | 0.26276315 | 0.2245022 | 0.68126557 | 0.83835842 |
| **HRD3** | 0.69348641 | 0.02420261 | 0.0612112 | 0.27473743 | 0.3749641 | 0.61220568 |
